# Anticipatory discovery of entry inhibitors against emerging viruses guided by viral phylogeny

**DOI:** 10.64898/2026.07.17.739150

**Authors:** Rodrigo Arce, Iván Andreu-Moreno, Jérémy Dufloo, Rafael Sanjuán

**Affiliations:** Institute for Integrative Systems Biology, Universitat de València - CSIC, Paterna, València, Spain

## Abstract

Emerging viruses pose a major threat to global health, underscoring the need to improve antiviral preparedness. Here, we developed a phylogenetically-informed framework to identify entry inhibitors with potential activity against yet-to-emerge viruses. First, using pseudotypes displaying the receptor-binding proteins (RBPs) of 8 enveloped RNA viruses from different families, we screened a library of 2,320 FDA-approved compounds to identify candidate entry inhibitors. Hits were tested for potency, selectivity, consistency across cell types and pseudotyping vectors and, in some cases, were validated using authentic viruses. Then, to define the breadth of antiviral activity, 25 selected drugs were assayed against an expanded panel of 68 RBPs from 13 families, as well as against pairs of closely related RBPs. This revealed both narrow and broad-range inhibitors, including selective estrogen receptor modulators, alkaloids, aminoquinolines, and anidulafungin, which showed particularly broad activity. Importantly, RBPs from the same phylogenetic cluster frequently displayed correlated drug-sensitivity profiles, indicating that antiviral effects are predictable across closely related viruses. Our findings provide a proof of concept for anticipatory antiviral discovery, showing that phylogenetic relationships can guide the identification of entry inhibitors against potential future zoonotic threats.

## Introduction

The series of viral outbreaks over the past two decades calls for effective measures to strengthen pandemic preparedness. These events include the emergence of Severe Acute Respiratory Syndrome (SARS) in 2002-2003, the global spread of the 2009 H1N1 influenza pandemic, the emergence of Middle East Respiratory Syndrome (MERS) in 2012, the West African Ebola epidemic (2014-2016), and the Zika epidemic (2015-2016) among others. More recently, SARS-CoV-2 generated unprecedented global socioeconomic and public-health impacts, and highly pathogenic avian influenza A (H5N1) has reemerged with widespread outbreaks among birds, marine mammals, and cows, as well as sporadic human cases. Additional emerging or re-emerging viruses include Oropouche virus, which has expanded throughout South American and Caribbean countries in 2023-2024 and Chikungunya, which has produced large, geographically diverse waves in 2024-2025 among other outbreaks.

Improved surveillance, diagnostics, vaccines and antivirals are thus required to increase our preparedness against the next pandemic caused by the so-called “Pathogen X” (Chiu et al., 2023; Mehand et al., 2018; Sikkema & Koopmans, 2025; S. Wang et al., 2024). The majority of emerging viruses have so far originated from mammal or avian reservoirs (Woolhouse et al., 2012), and enveloped RNA viruses tend to show the broadest host ranges and highest cross-species transmissibility (Olival et al., 2017; Valero-Rello & Sanjuán, 2022). Therefore, although it is currently not possible to ascertain the specific identity of the next viral pathogen, our efforts should be directed preferentially against enveloped RNA viruses infecting these groups of hosts. However, the vast majority of potentially human-infective viruses remain undiscovered, and culturing of many of the known viruses is highly restricted by technical or biosafety concerns.

To address these challenges, engineered viral systems have been implemented to reproducibly mimic some key aspects of viral infection cycles. Notably, pseudotypes are chimeric viral particles in which the receptor-binding protein (RBP) of a virus of interest is incorporated into a viral vector, typically lentiviruses or vesicular stomatitis virus (VSV). Pseudotyping has been applied to most families of enveloped RNA viruses and can faithfully reproduce RBP-driven processes such as receptor usage, cellular tropism, antibody-mediated neutralization, and sensitivity to entry inhibitors (Y. Wang et al., 2023). Therefore, pseudotypes provide a potent tool for targeting the RBPs of a wide variety of potentially emerging viruses. Recently, we developed a large panel of VSV and lentiviral pseudotypes carrying the RBPs of > 100 different RNA viruses belonging to 14 different families, and showed that most RBPs from mammalian and avian viruses are compatible with human cell entry factors, thus allowing to study a key aspect of human infectivity under controlled laboratory conditions (Dufloo et al., 2025).

Although vaccination is arguably the most efficient long-term control measure against infectious diseases, it also has limitations, since vaccine development and deployment can be outpaced by rapid viral emergence. Moreover, vaccination coverage needs to be extensive, which makes it sensitive to the growing problem of vaccine hesitancy. In addition, vaccines can be ineffective in immunocompromised individuals, including chronic patients, the elderly, and pregnant women. On the other hand, novel drug discovery remains slow and costly. This has led to explore drug repurposing as a tractable route to accelerate therapeutic responses, since candidate compounds already have known safety and pharmacokinetic profiles (Almulhim et al., 2025). This strategy has been followed for several viruses, mainly, SARS-CoV-2 (Riva et al., 2020; Touret et al., 2020; Xiong et al., 2021), Ebola and other filoviruses (Anantpadma et al., 2016; Cheng et al., 2015; Kouznetsova et al., 2014; Lee et al., 2018), and arenaviruses (Cao et al., 2021; Wan et al., 2021).

In this study, we screened a library of 2,320 FDA-approved compounds against a panel of pseudotypes bearing the RBPs from 8 viruses across different viral families to identify candidate entry inhibitors. Hits were verified using alternative pseudotyping vectors and cell lines. Next, we tested these compounds in an extended pseudotype panel of 60 RBPs belonging to 13 viral families to delineate their breadth of action. To more precisely ascertain whether viral phylogenetic clustering allows predicting the efficacy of entry inhibitors, we constructed and tested additional pseudotypes carrying RBPs from viruses closely related to compound-sensitive RBPs. Finally, selected authentic viruses were used to confirm the activity of these compounds. Our study establishes an evolutionary-guided approach to increase our preparedness against future emerging viruses.

## Materials and methods

### Cells

HEK-293T and Vero cells were cultured in Dulbecco’s modified Eagle medium (DMEM) supplemented with 10% fetal bovine serum (FBS), 10 U/ml penicillin, 10 µg/ml streptomycin (Gibco), 250 ng/ml amphotericin B (Gibco), and 1% non-essential amino acids (NEAA; Gibco). A549, OVCAR-8, and SNB-19 cells were cultured in RPMI (Gibco) supplemented with 10% FBS, 10 U/ml penicillin, 10 µg/ml streptomycin (Gibco), 250 ng/ml amphotericin B (Gibco), and 1% non-essential amino acids (NEAA; Gibco). HUVECs cells were provided by Prof. Isabel Fariñas (Universitat de València, Spain) and cultured in Endothelial Cell Growth Medium (PromoCell) supplemented with 18.5% Growth Medium Supplement Mix (PromoCell), 10 units per ml penicillin, 10 µg/ml streptomycin (Gibco) and 250 ng/ml amphotericin B (Gibco). All cells were maintained at 37°C in 5% CO_2_. Cells were regularly shown to be free of mycoplasma contamination by PCR.

### VSV pseudotype production

VSV pseudotypes were produced as detailed in our previous work (Dufloo et al., 2025) with minor modifications. Briefly, T75 culture flasks were pre-coated with poly-D-lysine (Gibco) for 2 h at 37 °C, rinsed with sterile water, and subsequently seeded with 8 × 10⁶ HEK-293T cells. After 24 h, cells were transfected with 18 µg of pcDNA3.1 vector encoding the viral RBP, using PolyJet (SignaGen) according to the manufacturer’s protocol. For negative controls, an empty pcDNA3.1 vector was transfected in parallel (Empty control). At 24 h post-transfection, cells were infected at a multiplicity of infection (MOI) of 3 infectious units per cell for 1 h at 37 °C with a VSV lacking the glycoprotein gene G, encoding GFP (VSVΔG-GFP), and previously pseudotyped with its own RBP, VSV-G. Following infection, cells were washed three times with phosphate-buffered saline (PBS) and replenished with 8 mL of DMEM supplemented with 2% FBS. After 24 h, supernatants were harvested, clarified by centrifugation at 2,000 × g for 10 min, filtered through a 0.45 µm membrane, aliquoted, and stored at −80 °C until use. The RBP panel described in previous work (Dufloo et al., 2025) was expanded to include the RBPs of La Crosse virus (obtained from pHCMV_LACV_GP, a gift from Dr. Feng Zhang (Massachusetts Institute of Technology, USA); Addgene plasmid # 207286; RRID: Addgene_207286), Oropouche virus (de novo synthetized, IDT), Aransas Bay virus, Sudan virus, Hunter Island virus, Machupo virus, and Aruac virus (de novo synthetized, Genscript), as well as Ebola virus (Kikwit strain) and Bundibugyo (provided by Dr. Rafael Delgado-Vázquez, Instituto de Salud Carlos III, Spain).

### Lentivirus pseudotype production

HIV-based pseudotypes were produced as previously described by our group (Dufloo et al., 2025). Instead of using pCMVΔR8.2 as the packaging plasmid, we used psPAX2. psPAX2 was provided by Dr. Didier Trono (École polytechnique fédérale de Lausanne, Switzerland); Addgene plasmid # 12260; RRID:Addgene_12260). Briefly, six-well culture plates were coated with poly-D-lysine (Gibco) for 2 h at 37 °C, washed with sterile water, and seeded with 1×10⁶ HEK-293T cells. After 24 h, cells were transfected with 0.83 µg each of psPAX2 packaging plasmid, pTRIP-GFP, and the plasmid encoding the viral RBP, using Lipofectamine 3000 (Invitrogen) according to the manufacturer’s protocol. Supernatants were harvested 48 h post-transfection, clarified by centrifugation at 2,000 × g for 10 min, aliquoted, and stored at −80 °C until further use. The pseudotype with the RBP of Junin virus was harvested 72 h post-transfection. Lentiviral pseudotypes were titrated in HEK-293T cells.

### Screening (Phase 1)

The DiscoveryProbe FDA-Approved Drug Library (APExBIO, Catalog No. L1021), which contains 2,320 FDA-approved compounds, was screened against a panel of 8 viral pseudotypes, each carrying the RBP of a virus from a different family: Sindbis virus (SINV, *Togaviridae)*, Thogoto virus (THOV, *Orthomyxoviridae*), Leopards Hill virus (LPHV, *Nairoviridae*), Rabies virus (RABV, *Rhabdoviridae*), Sin Nombre virus (SNV, *Hantaviridae*), Junin virus (JUNV, *Arenaviridae*), Ebola virus Makona strain (EBOV, *Filoviridae*), and La Crosse virus (LACV, *Peribunyaviridae*). For the assay, 15,000 HEK-293T cells per well were seeded in 96-well plates in 50 µL of complete DMEM. The following day, 25 µL of each compound diluted to 40 µM was added to each well. After a 4 h preincubation, 25 µL of pseudotype (100-1500 infectious units) was added, resulting in a final compound concentration of 10 µM. At 24 h post-inoculation, cells were imaged using the Incucyte SX5 Live-Cell Analysis System (Sartorius), and GFP-positive cells were quantified automatically with the Incucyte Analysis software (v2024BRev2). Each 96-well plate included eight untreated wells that served as negative controls. Infection rates were calculated by dividing the GFP count in compound-treated wells by the mean GFP count in untreated controls. A total of 100 candidate compounds were identified according to the following criteria: (i) infection as determined by GFP signal decreased by > 90% relative to untreated controls; (ii) for these compounds, if the cell monolayer showed signs of cytotoxicity, they were advanced to cytotoxicity testing only for confirmation and were not further evaluated for inhibition; (iii) compounds with 70-90% infection reduction were considered for further evaluation if they targeted viruses with few or no other candidate inhibitors. Data from Phase 1 are available in Supplementary Data S1.

### Antiviral and cytotoxicity assays (Phase 2)

Dose-response inhibition (IC) and cytotoxicity (CC) assays were performed by treating cells with serial dilutions of the compounds. For the first set of 10 compounds, the concentrations tested were 40, 20, 10, 2.5, 0.62, 0.16, 0.04, and 0.01 µM. Since the lowest concentrations produced minimal effects, subsequent assays were performed using the range 40, 20, 10, 5, 2.5, 1.25, 0.62, and 0.31 µM. We discarded 19 compounds in the early stages of Phase 2 due to obvious cytotoxic effects (high CC_50_) or because the compounds were reported to targeted post-entry stages and hence inhibited VSV replication. IC assays were performed for the remaining 81 compounds against the pseudotypes that showed inhibition in Phase 1. IC₅₀ and CC₅₀ values were determined using nonlinear regression to a logistic model with GraphPad Prism 10.6.0. In this phase, compounds were selected based on the following criteria: (i) specificity index (SI) > 10, calculated as CC₅₀/IC₅₀ for at least one virus; (ii) IC₅₀ < 6 µM; (iii) CC₅₀ > 20 µM; and (iv) cell viability at 10 µM > 85%. Since most of the initial hits targeted EBOV and JUNV pseudotypes, additional criteria were applied to retain only a subset of the compounds active against these two pseudotypes and ensure broader viral coverage: (v) compounds active against more than one virus were prioritized over those with similar efficacy against a single virus; (vi) a maximum of 10 compounds exclusively active against a single virus were retained, prioritizing those that were not previously reported in the literature; (vii) for viruses with only one or two inhibitors, thresholds were relaxed. Data from Phase 2 are available in Supplementary Data S2.

### Orthogonal assays (Phase 3)

This phase was carried out to evaluate cell type and pseudotype vector specificity. First, compounds were tested against VSV pseudotypes at 40, 10, 4, and 0.4 µM in five different cell lines (A549, OVCAR-8, SNB-19, HUVEC, and HEK-293T for confirmation) to quantify both cytotoxicity and inhibition. Second, compounds were tested at 10 µM in HEK-293T cells using both VSV- and lentivirus-based pseudotypes. Compounds were selected based on the following criteria: (i) cells yielding fewer than 25 counts were considered non-permissive (e.g., Junin, Ebola, Leopards Hill, and Sindbis did not infect HUVEC); (ii) a compound was considered effective if it reduced infection with VSV-based pseudotypes to < 25% of mock-infected levels in at least (n−1) out of n permissive cell lines at 10 µM; (iii) cell viability had to remain ≥ 85% in at least 4 of the 5 tested cell lines at 10 µM; and (iv) compounds were discarded if infection levels for lentiviral pseudotypes remained > 30% of mock after treatment at 10 µM. Data from Phase 3 are available in Supplementary Data S3.

### Compound screening against an extended panel of viral pseudotypes (Phase 4)

Compounds retained in Phase 3 were used to assess their efficacy in an extended pseudotype panel. In this phase, it was necessary to treat pseudotypes with anti-VSV-G monoclonal antibody, since RBPs showing lower pseudotyping efficacy may yield significant background from the carryover of VSV-G used during their production. Therefore, we first verified that all except two of the compounds retained in Phase 3 showed > 90% inhibition at 10 µM in the presence of an anti-VSV-G monoclonal antibody. Next, HEK-293T cells (15,000 cells/well) were seeded in 96-well plates with 50 µL of complete DMEM and incubated overnight. The following day, 25 µL of each test compound, diluted to 40 µM, was added to the wells and incubated for 4 h. During the treatment, equal volumes of VSV pseudotypes and anti-VSV-G monoclonal antibody were mixed for 30 mins at 37 °C. Subsequently, 25 µL of each pseudotype-antibody mix (25-1500 infectious units, depending on the pseudotype) was added, resulting in a final compound concentration of 10 µM. At 24 h post-inoculation, cells were imaged using the Incucyte SX5 Live-Cell Analysis System (Sartorius), and GFP-positive cells were quantified automatically with the Incucyte Analysis software (v2024BRev2). Each 96-well plate contained 6 negative controls: four wells with DMEM only and two wells infected with an empty VSV pseudotype carrying no RBP (VSV-Empty). In addition, two non-treated 96-well plates were included in each experimental run and used to calculate infection rates. All experiments were performed in duplicate. To minimize the impact of handling errors, a third replicate was carried out in the following situations: (i) the fold-change reduction in infectivity between replicates diverged by more than twofold, except when treated wells contained fewer than 10 infectious units and/or the mean fold change was below 0.05; (ii) empty controls in any non-treated plate exceeded three counts, or when their mean value was greater than one; and (iii) DMEM-only control well showed more than one count. To preserve the duplicate structure of the dataset, the most deviating value among the three replicates was discarded. Data from Phase 4 are available in Supplementary Data S4.

### Authentic virus assays

Wells were coated with poly-D-lysine (Gibco) for 30 min at 37°C, then washed with water. The cells were seeded to reach 80-90% confluence on the day of the assay. Cells were treated with the compounds for 4 h, inoculated for 1 h and washed 3 times with PBS. All assays with full-viruses were performed in HEK-293T cells. Infections with THOV isolate SiAr126 were performed at MOI of 0.01 plaque forming units (PFU) per cell. Viral production was titrated at 48 h post inoculation (hpi) by plaque assay in Vero cells. For lymphocytic choriomeningitis virus (LCMV, *Arenaviridae*), infections were performed at MOI of 0.1 PFU/cell and viral growth was quantified by RT-qPCR at 48 hpi using previously described primers (McCausland & Crotty, 2008). Infections with a GFP-expressing SINV were performed at MOI of 0.1 FFU/cell and viral growth was evaluated at 16 hpi by the foci formation assay in HEK-293T cells. LCMV and SINV-GFP were provided by Drs. Celia Perales (Centro de Biología Molecular Severo Ochoa, Spain) and Ron Geller (I2SysBio, Spain), respectively. Data from infections with authentic viruses are available in Supplementary Data S5.

### Hierarchical clustering of antiviral effects

A matrix of normalized inhibition values was used. Pairwise distances between viruses were computed using the correlation distance 1 – r, where r is the Pearson correlation, as this metric emphasizes similarity in inhibition profiles regardless of magnitude. Hierarchical clustering was performed using the average linkage method (hclust function in R). Clustering was computed for RBPs and for compounds. The resulting dendograms were integrated into a heatmap representation of the inhibition matrix. All analyses were implemented in R (v.4.3.2) using the packages pheatmap, dendextend, RColorBrewer, mclust, cluster, svglite, and vegan.

### Classification of antiviral hits

Primary antiviral hits identified for each viral system were annotated according to their mechanism-of-action pathway using the classification provided by the compound library supplier. The distribution of pathway categories across viral systems was visualized as pie charts using R (v4.3.2) with the ggplot2 package (Fig. S1). Full annotation details are provided in Supplementary Data S5.

### Sequence analysis

RBP phylogenies were constructed from amino acid sequences by carrying out alignments with MAFFT and inferring maximum likelihood trees with IQ-TREE with automatic model selection and 1,000 ultrafast bootstrap replicates. Due to the diversity of viral families analyzed, trees were midpoint-rooted for visualization purposes. Trees were visualized and annotated using iTOL. To compare pairs of RBP sequences, their amino acid sequence was calculated using pairwise alignments performed with the EMBOSS Needle tool available through the EMBL-EBI server. Alignments were generated using the Needleman-Wunsch algorithm with default parameters (BLOSUM62 substitution matrix, gap opening penalty of 10, and gap extension penalty of 0.5). Percentage identity was defined as the number of identical residues divided by the total alignment length, including gaps.

## Results

### Screening and validation

We established a screening workflow using VSV-based pseudotypes bearing the RBPs of viruses from 8 different families: Sindbis virus (SINV, *Togaviridae)*, Thogoto virus (THOV, *Orthomyxoviridae*), Leopards Hill virus (LPHV, *Nairoviridae*), Rabies virus (RABV, *Rhabdoviridae*), Sin Nombre virus (SNV, *Hantaviridae*), Junin virus (JUNV, *Arenaviridae*), Ebola virus Makona strain (EBOV, *Filoviridae*), and La Crosse virus (LACV, *Peribunyaviridae*) (Fig. 1a). An initial screen of 2,320 FDA-approved compounds (Phase 1) identified 100 potential hits that reduced viral infectivity by > 90% (Fig. 1b; Supplementary Data S1). Notably, the majority of these candidate hits inhibited the entry of EBOV (64) and JUNV (23) (Fig. S1). In Phase 2, candidate hits were subjected to dose-response characterization to determine CC_50_ and IC_50_ values and calculate the selectivity index (SI; Fig. 1c) and we identified 81 compounds with SI > 10 for all viruses except RABV (Supplementary Data S2; Figs. S2-S6). Following these criteria, 32 drugs were advanced to Phase 3 for orthogonal validation (Supplementary Data S3). This stage assessed cell-type specificity across five cell types (HEK-293T, A549, OVCAR-8, SNB-19, and HUVEC primary cells), revealing significant cytotoxicity in HUVECs for 15 candidates and excluding four due to cell-line-dependent effects (Fig. 1d). To minimize system-specific artifacts, candidates were further validated using lentivirus-based pseudotypes in HEK-293T cells. This step excluded two additional drugs that showed reduced inhibition using the lentiviral backbone compared to the VSV system (Fig. 1e). Ultimately, after discarding two additional candidates due to possible interference between the anti-VSV-G monoclonal antibody and compound activity, we retained 24 drugs that inhibited one or multiple pseudotypes, totaling 50 RBP-compound active pairs. No candidates were retained at this stage for LACV and RABV. After a literature search, we included an additional compound, GRP-60367, reported to be active against RABV (Pont et al., 2020) (Supplementary Data S2).

**Figure 1.**
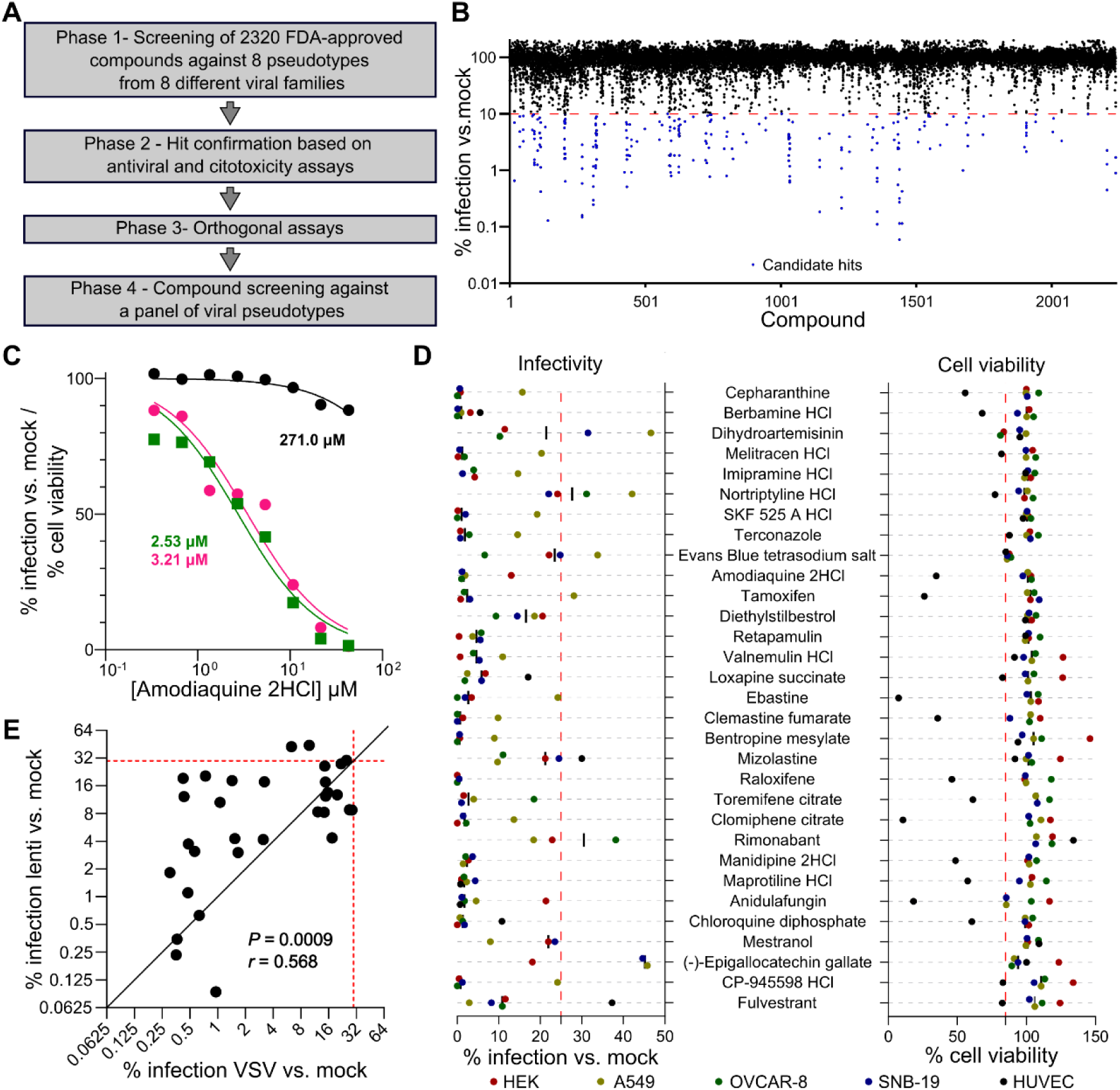
Pipeline and result summary for the screening of FDA-approved compounds against viral pseudotypes. (A) Schematic overview of the four-phase screening process. (B) Phase 1: Screening results showing percent infection relative to untreated controls. Each dot represents a drug-RBP assay, and blue dots denote compounds selected as candidate hits. (C) Phase 2: Example of dose–response curves for Amodiaquine 2HCl showing antiviral activity against THOV (green) and EBOV (pink) pseudotypes, as well as the corresponding cell viability (black). IC_50_ or CC_50_ data are shown with their respective colors. (D) Phase 3-1: Infectivity (left) and cell viability (right) of candidate compounds tested in HEK293T, A549, OVCAR-8, SNB-19, and HUVEC cells at 10 µM. The red dashed lines indicate 25% infection cutoff and 85% viability threshold. (E) Phase 3-2: Comparison of antiviral activity for compounds tested in both VSV- and lentivirus-based pseudotypes. The red dashed line indicates the 30% infection cutoff. The black line indicates a slope of 1. The Pearson correlation coefficient (r) and p-value (P) for the log_2_ transformed percentage of infection are indicated in the graph.

### Association between drug activity and viral taxonomy across 68 RBPs

We sought to characterize the inhibitory breadth of the 25 drugs at 10 µM across 68 pseudotypes carrying RBPs from 13 distinct viral families. Overall, we identified 218 RBP–drug combinations showing >80% inhibition. We then calculated the frequency with which each drug reduced viral infection by at least 80% among RBPs belonging to the same viral family (Fig. 2). The data revealed distinct family-specific susceptibility patterns. Notably, *Filoviridae* RBPs showed high sensitivity to alkaloids (cepharanthine and berbamine) and aminoquinolines (amodiaquine and chloroquine), suggesting shared entry mechanisms within this family. Additionally, selective estrogen receptor modulators (SERMs), including tamoxifen, toremifene, and clomiphene, as well as the antihistamine ebastine, affected *Filoviridae* and *Arenaviridae* RBPs with similar efficiency. Anidulafungin emerged as the most broadly active inhibitor in our library (41.1%), effectively targeting not only these families but also members of the order Bunyavirales (*Nairoviridae*, *Phenuiviridae*, and *Peribunyaviridae*) as well as *Orthomyxoviridae* RBPs. This broad spectrum of activity suggests disruption of conserved entry pathways.

**Figure 2.**
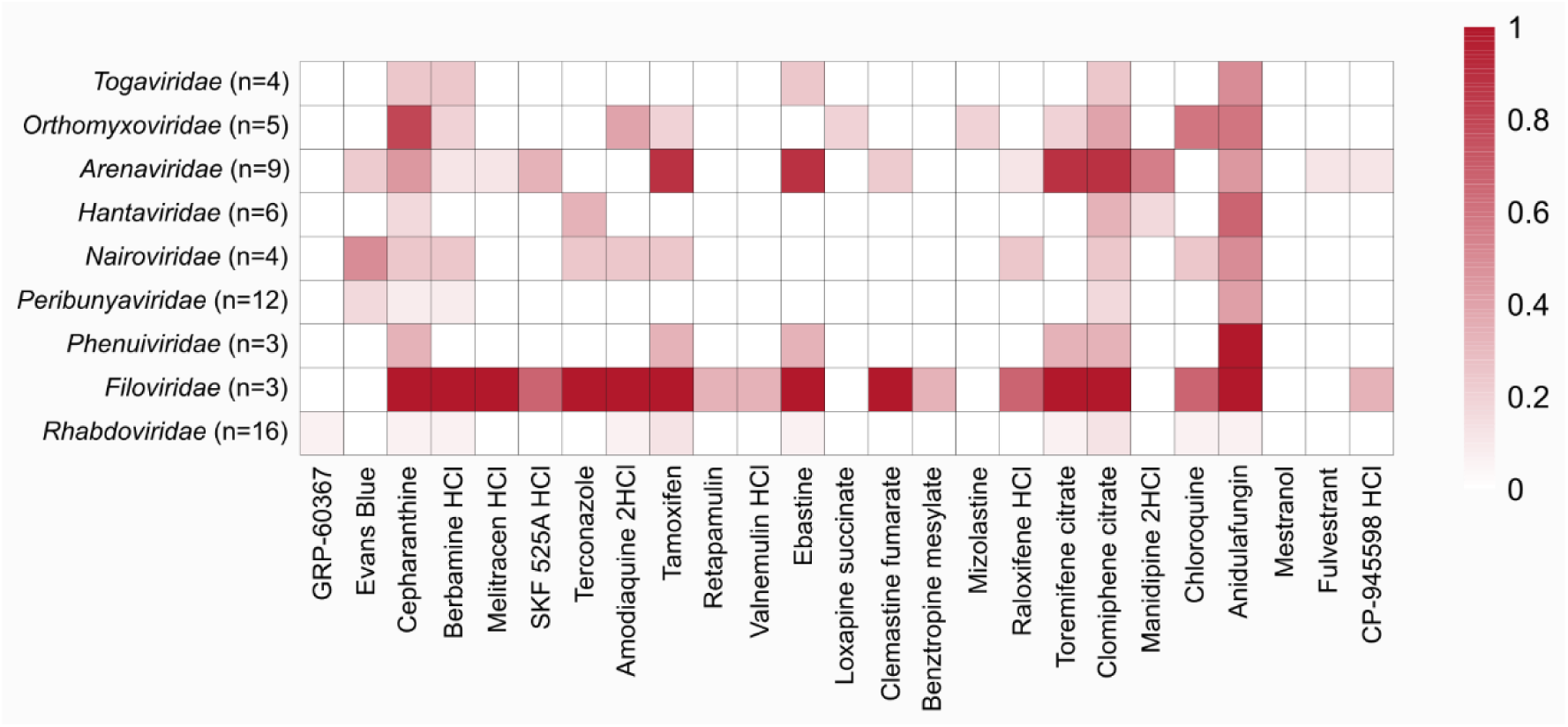
Family-specific activity of selected drugs. Heatmap showing the frequency of RBPs within each family (n indicated in parentheses) showing > 80% inhibition at 10 µM with each selected compound. The color scale ranges from 0 (white, no virus within the family inhibited) to 1 (dark red, 100% of viruses within the family inhibited). Families with two or fewer members are not shown.

To systematically classify viral inhibition profiles, we performed hierarchical clustering analysis (Fig. 3). This revealed two major clades of compounds with consistent antiviral activity, broadly separating SERMs and ebastine from alkaloids and aminoquinolines. These patterns suggest the presence of shared inhibitory mechanisms across phylogenetically distant viral taxa. Viral clustering partially mirrored viral taxonomy, although more complex relationships also emerged. While *Arenaviridae* RBPs generally clustered together (8/10), they also grouped closely with Itacaiunas virus (ITAV; *Rhabdoviridae*), indicating similar susceptibility profiles despite substantial phylogenetic divergence. Within the *Rhabdoviridae*, nine pseudotypes formed a distinct clade characterized by low overall inhibition, whereas RABV and Gannoruwa bat lyssavirus (GBLV) clustered separately, likely due to their sensitivity to the control inhibitor GRP-60367. Additional clusters were observed for members of the *Peribunyaviridae* and *Orthomyxoviridae*, excluding THOV.

**Figure 3.**
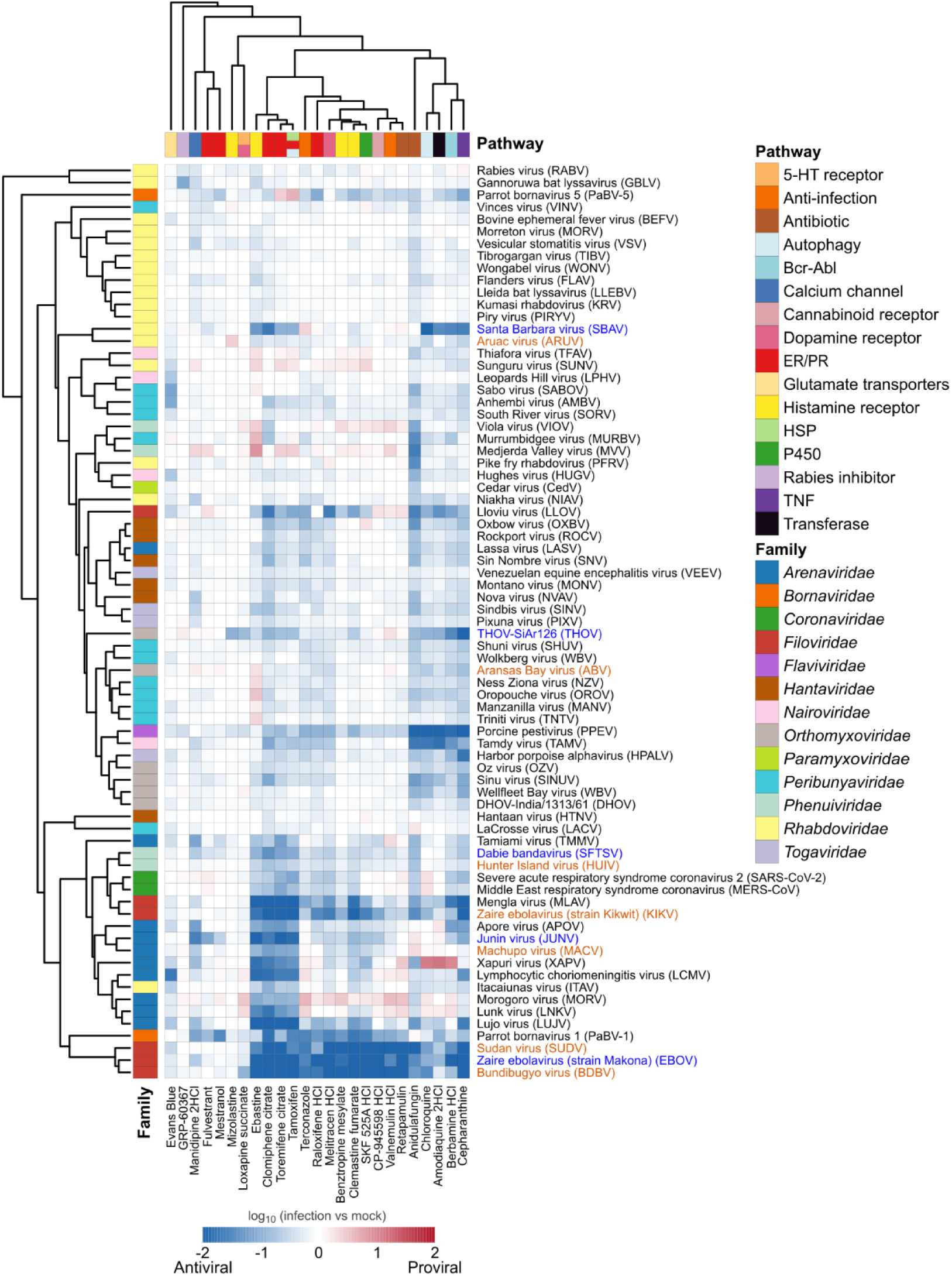
Hierarchical clustering of antiviral activities of FDA-approved compounds across enveloped RNA viruses. The heatmap depicts the log_10_ fold change in infection relative to untreated controls, with blue indicating inhibition and red indicating proviral activity. Both rows and columns were hierarchically clustered using the correlation distance and the average linkage method to reveal similarity patterns among inhibition profiles. Colored bars on the left denote viral families, and colors at the top indicate the pathways targeted by each compound. Blue and orange virus names represent pairs used to test for similarities between related RBPs (blue: present in the initial panel; orange: newly synthesized RBP; see text). Pathway: ER/PR (estrogen/progesterone receptor); HSP (heat shock protein); TNF (tumor necrosis factor).

### Drug sensitivity is conserved among related RBPs

The above results reveal that RBP drug sensitivities show only some level of clustering according to viral family. To more precisely examine whether the effect of a compound on a given RBP can be extrapolated to related viruses, we build RBP phylogenies for 5 different families (Figs. S7-S11). New RBPs were selected as phylogenetic neighbors of assayed RBPs that showed strong drug sensitivity (more than 7 compounds with < 20% infection vs. control) in the screening with the 68 pseudotypes. The new RBPs belonged to the same viral genus and clustered within the same well-supported phylogenetic clade (bootstrap ≥80) as the assayed RBPs used as reference. The selected RBP pairs are shown in Table 1.

**Table 1.** Reference RBPs and newly selected RBPs based on phylogenetic relatedness. Sequence identity indicates the percentage of amino acid identity between RBPs.

|  |  |  |  |  |  |  |  |
| --- | --- | --- | --- | --- | --- | --- | --- |
| Reference virus | SFTSV | THOV | JUNV | SBAV | EBOV | EBOV | EBOV |
| New virus | Hunter Island virus (HUIV) | Aransas Bay virus (ABV) | Machupo virus (MACV) | Aruac virus (ARUV) | Ebola virus Kikwit strain (KIKV) | Bundibugyo virus (BDBV) | Sudan virus (SUDV) |
| Family | Phenui | Orthomyxo | Arena | Rhabdo | Filo | Filo | Filo |
| Genus | Bandavirus | Thogotovirus | New World clade B | Arurhavirus | Ebolavirus | Ebolavirus | Ebolavirus |
| Sequence identity (%) | 54.5 | 42.1 | 69.2 | 23.3 | 97.2 | 64.9 | 54.7 |
| Phylogeny | Fig. S7 | Fig. S8 | Fig. S9 | Fig. S10 | Fig. S11 | Fig. S11 | Fig. S11 |

Each of these 7 new RBPs were synthesized, used to produce VSV pseudotypes, and tested with the 25 selected compounds. Overall, sensitivity profiles were strongly correlated for each pair of related RBPs (Fig. 4). This was particularly evident for the *Filoviridae*, where the response of the EBOV pseudotype demonstrated a high correlation with those of the KIKV (r = 0.897), BDBV (r = 0.912), and SUDV (r = 0.862) pseudotypes. Similar patterns were observed in the *Phenuiviridae* (HUIV vs. SFTSV, r = 0.931) and *Arenaviridae* (MACV vs. JUNV, r = 0.754). Even for RBPs that showed lower sensitivity to the selected compounds, such as THOV and SBAV, the correlations remained significant (r = 0.700 and r = 0.652, respectively; *P* < 0.01), indicating shared entry vulnerabilities. Importantly, 5 out of the 7 new RBPs clustered together with their reference RBPs when included in the hierarchical clustering analysis (Fig. 3), KIKV and ABV being the exceptions. While KIKV clustered with Mengla virus, a filovirus from another genus, ABV cluster in a clade of viruses from the *Peribunyaviridae* family. Overall, these results suggest that the entry inhibitor sensitivities of newly emerging viruses may be predicted from those of known viruses belonging to the same phylogenetic cluster, even if their RBPs have diverged substantially.

**Figure 4.**
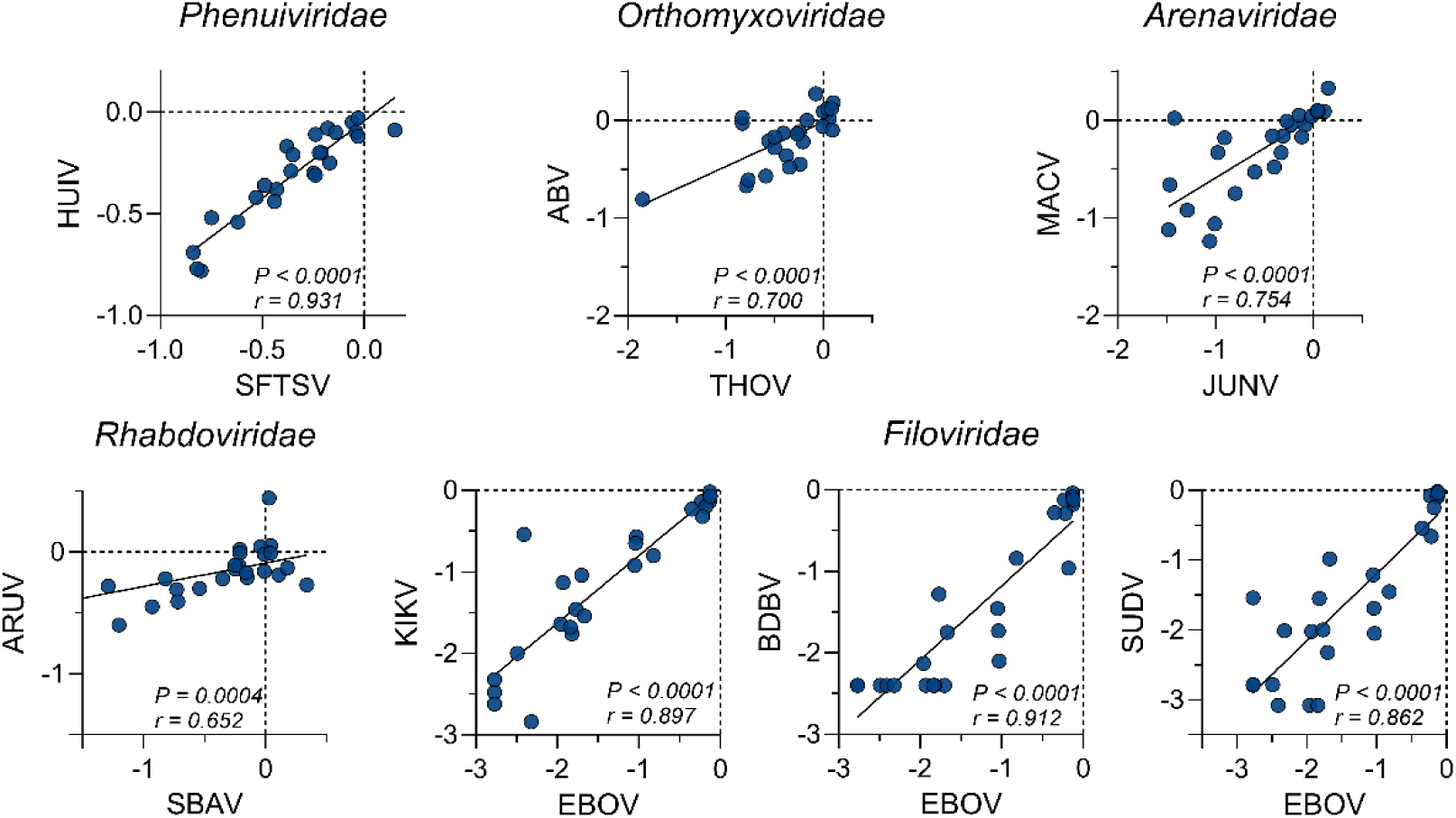
Phylogenetically related RBPs show correlated responses to antiviral compounds. Scatter plots showing the correlation between the compound inhibition profiles of pseudotypes carrying related RBPs. Each dot represents one of the 25 selected compounds. Solid lines show the least-squares linear regression. The Pearson correlation coefficient (r), and p-value (*P*) are indicated within each panel. Data are expressed as the log_10_ fold change in infection relative to untreated controls. Dashed lines mark the zero-effect threshold. HUIV: Hunter Island virus; SFTSV: Severe fever with thrombocytopenia syndrome virus; ABV: Aransas Bay virus; MACV: Machupo virus; ARUV: Aruac virus; SBAV: Santa Barbara virus; KIKV: Ebola virus; BDBV: Bundibugyo virus; SUDV: Sudan virus.

### Validation with authentic viruses

To test whether the entry hits identified with pseudotypes were active against authentic viruses, we selected 8 compounds for THOV, 3 for LCMV, and 1 for SINV-GFP. The authentic viruses were selected based on availability and biosafety (BSL-2) criteria. The results confirmed the antiviral activity of each compound (Fig. 5). For THOV, amodiaquine and berbamine stood out as the most potent inhibitors, driving viral titers down by 3 to 4 orders of magnitude (one-way ANOVA with post hoc Dunnet’s test: *P* < 0.01). The LCMV titer was reduced > 30-fold when cells were treated with toremifene, tamoxifen, or evans blue (one-way ANOVA with post hoc Dunnet’s test: *P* < 0.001). Finally, clomiphene reduced the SINV-GFP titer by approximately two orders of magnitude (two-tailed t-test; *P* < 0.001), as previously seen by others but not associated to viral entry (Mudgal et al., 2022). Hence, the antiviral effects identified with pseudotypes were reproduced with authentic viruses.

**Figure 5.**
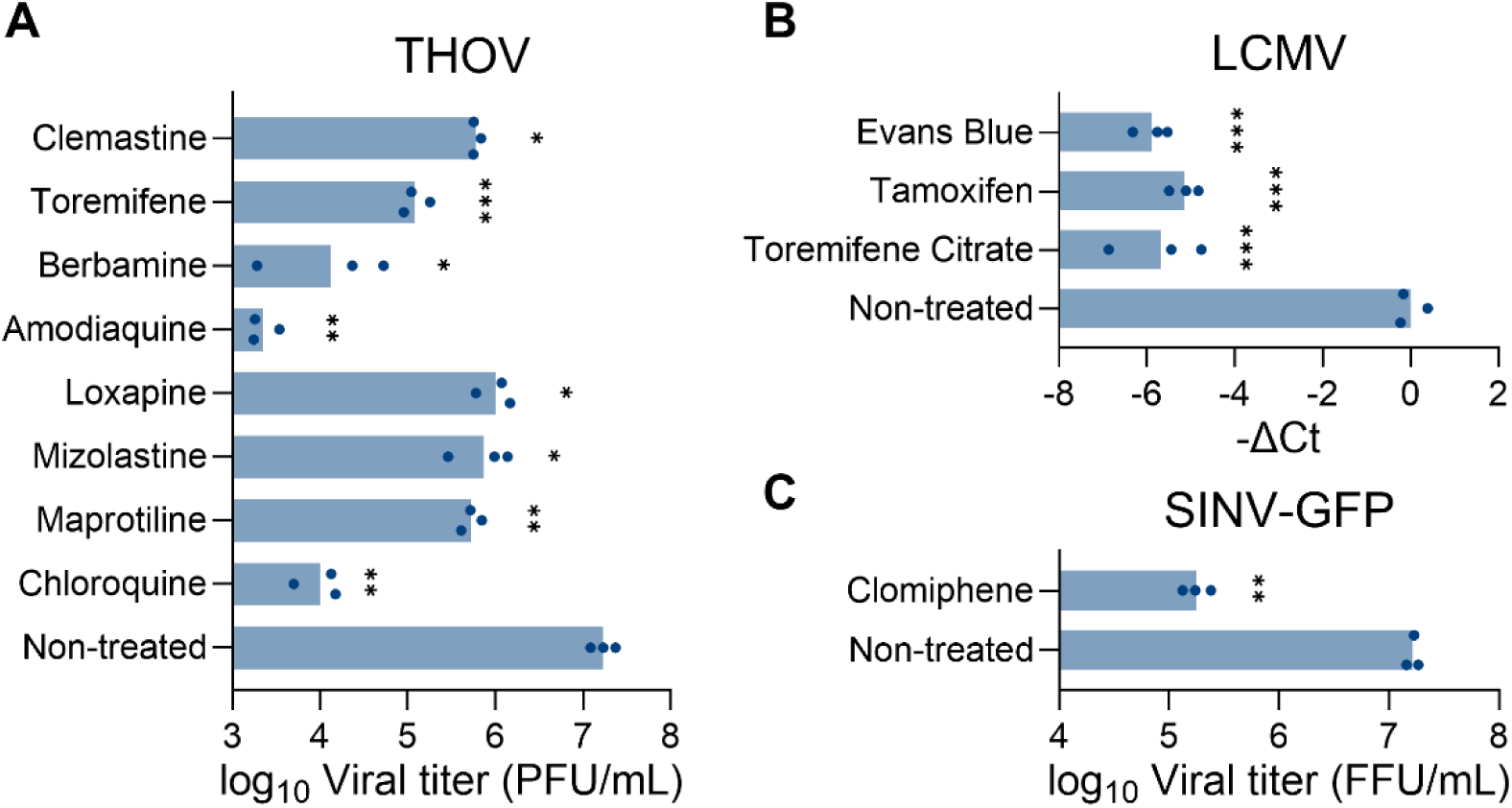
Assays with authentic viruses. (A) Inhibition of THOV. Titer is expressed as plaque former units (PFU) per mL. (B) Inhibition of LCMV. Viral growth was quantified by RT-qPCR. (C) Inhibition of SINV-GFP. Titers were expressed as foci forming units (FFU) per mL. For THOV and LCMV the results were compared to non-treated controls using a one-way ANOVA with Dunnet correction, and for SINV-GFP a two-tailed t-test was performed. \*\*\**P* < 0.001; \*\**P* < 0.01; \**P* < 0.05.

## Discussion

Our results identify 218 active RBP-drug pairs showing > 80% inhibition of viral entry at 10 µM, the vast majority of which have not been previously described. The most effective compounds were the SERMs tamoxifen, toremifene, and clomiphene, which showed ≥90% inhibition of viral entry across 13, 13, and 16 pseudotypes, respectively. Similarly, anidulafungin and cepharanthine inhibited 14 pseudotypes with comparable efficacy to the SERMs. Importantly, these results were consistent across five cell lines and in both VSV- and lentivirus-based pseudotyping systems, supporting the robustness of the observed entry inhibition. Furthermore, the selected compounds also showed antiviral activity against three authentic viruses from different viral families. Consistent with these observations, some of the RBP-drug pair identified here have previously been reported to inhibit infection by authentic viruses in independent studies, including alphaviruses, EBOV, and SARS-CoV-2 supporting the biological relevance of the antiviral activity detected in our screening(Mudgal et al., 2022; Sun et al., 2017; Uematsu et al., 2025; Zu et al., 2021). Since the effects of these drugs were not uniform across pseudotypes carrying different RBPs and were consistent across two different pseudotyping systems and, the authentic viruses, RBPs are their only possible target and, therefore, these compounds may be classified as entry inhibitors.

By combining functional screening with comparative sequence analysis, we observe that patterns of inhibition tend to cluster according to phylogenetic relationships, indicating that entry vulnerabilities are, to some extent, conserved. Building on these observations, our results suggest a framework for pandemic preparedness based on mapping the activity of entry inhibitors onto viral phylogenies. By systematically profiling compound effects across related viruses, this approach could define susceptibility patterns that help identify candidate inhibitors for newly identified pathogens, including a potential “Pathogen X”, using only RBP sequence or structural information. In this context, a key question is the level of phylogenetic relatedness required for such extrapolations to be reliable. Our data indicate that classification at the family level is unlikely to be sufficient, and that finer resolution, such as relationships at the genus level or below are more informative. Nevertheless, we found similar drug sensitivity profiles even between RBPs different up to 50% in sequence. We attempted to correlate inhibition profiles with amino acid sequence identity among RBPs in order to define a lower threshold predictive of similar responses, but no clear patterns were identified. Defining these thresholds will be important to establish when antiviral activity can be meaningfully inferred for emerging viruses and to support the assembly of a pre-characterized repertoire of entry inhibitors.

Beyond its potential application for identifying entry inhibitors, this approach may also provide a useful framework to investigate shared and distinct entry pathways across viruses. By comparing inhibition profiles across related or even unrelated viruses, it becomes possible to infer commonalities in entry mechanisms as well as subtle functional differences. For instance, despite the close relatedness of EBOV strains, the Kikwit strain displays an inhibition profile more similar to Mengla virus than to the Makona strain, suggesting that relatively minor differences in their glycoproteins may translate into measurable functional divergence. In contrast, other ebolaviruses, such as Bundibugyo and Sudan virus, show profiles more closely aligned with Makona, supporting a more conserved entry phenotype within that group. These observations are consistent with previous work showing that limited sequence variation in viral glycoproteins can have disproportionate effects on receptor usage or entry efficiency (Fels et al., 2021; Negrete et al., 2007; Ruedas et al., 2018). In addition, the positioning of the ABV orthomyxovirus within a cluster of peribunyaviruses in our analyses (Fig. 3) is notable given that ABV was initially described as a bunyavirus based on the virion physicochemical properties (Briese et al., 2014; Yunker et al., 1979).

Several limitations of this study should be considered. First, most of our analyses rely on pseudotypes. To address this, we first used two orthogonal pseudotyping systems to avoid finding system-specific hits and then we validated a subset of our findings with authentic viruses, obtaining consistent results that support the relevance of the observed entry inhibition patterns. However, extending this validation across a broader range of viruses will be important to further assess the generality of our approach. In this context, it should be noted that many of the viruses that would be most informative to test are restricted to high-containment settings (BSL-3 or BSL-4), which currently limits systematic evaluation. These limitations make pseudotypes particularly useful for our proposed workflow. Second, while our results reveal a relationship between phylogenetic relatedness and compound activity, the extent to which these patterns can be generalized remains to be fully defined. Finally, the classification of compounds into broad, family-level, or more specific activity profiles should be interpreted within the scope of the viral panel analyzed here, and may evolve as additional viruses are incorporated.

Taken together, our results highlight the need for broader and more systematic efforts to identify antiviral entry inhibitors, either by expanding the chemical space explored or by incorporating a wider diversity of RBPs. Elucidating the mechanisms of action of the identified compounds will be important to support their translation into *in vivo* models, but screening of entry inhibitors in cell cultures using pseudotypes is nevertheless a powerful tool for the rapid identification of entry inhibitors in the context of viral emergence. The approach proposed here may offer a robust and scalable framework to support preparedness against future viral threats.

## Acknowledgements

We thank all the Virus Evolution Unit members for helpful comments. We thank J. Buigues Bisquert and R. Martínez-Recio for technical assistance.

## Author contributions

R.A., J.D. and R.S. designed the research; R.A. performed the research; R.A., J.D., I.A.- M. and R.S. analysed data; R.A. and R.S. wrote the paper; R.S. provided funding. All authors read and approved the final version of the manuscript.

## Supplementary Figures

**Figure S1.**
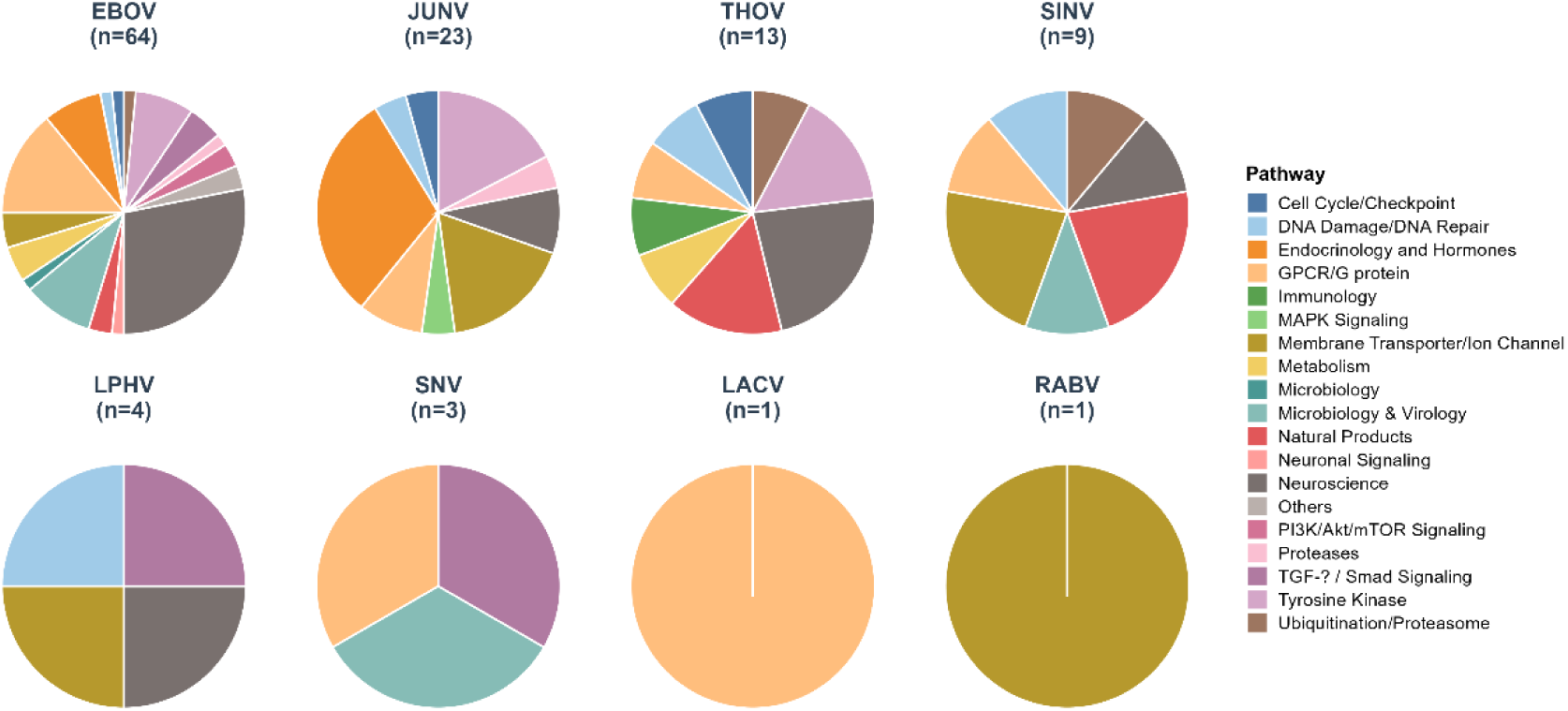
Pathway distribution of hits identified for each RBP. Pie charts represent the functional classification of compounds for each virus based on the annotation provided by APExBIO. For each viral pseudotype, the total number of selected hits is indicated in parentheses (n).

**Figure S2.**
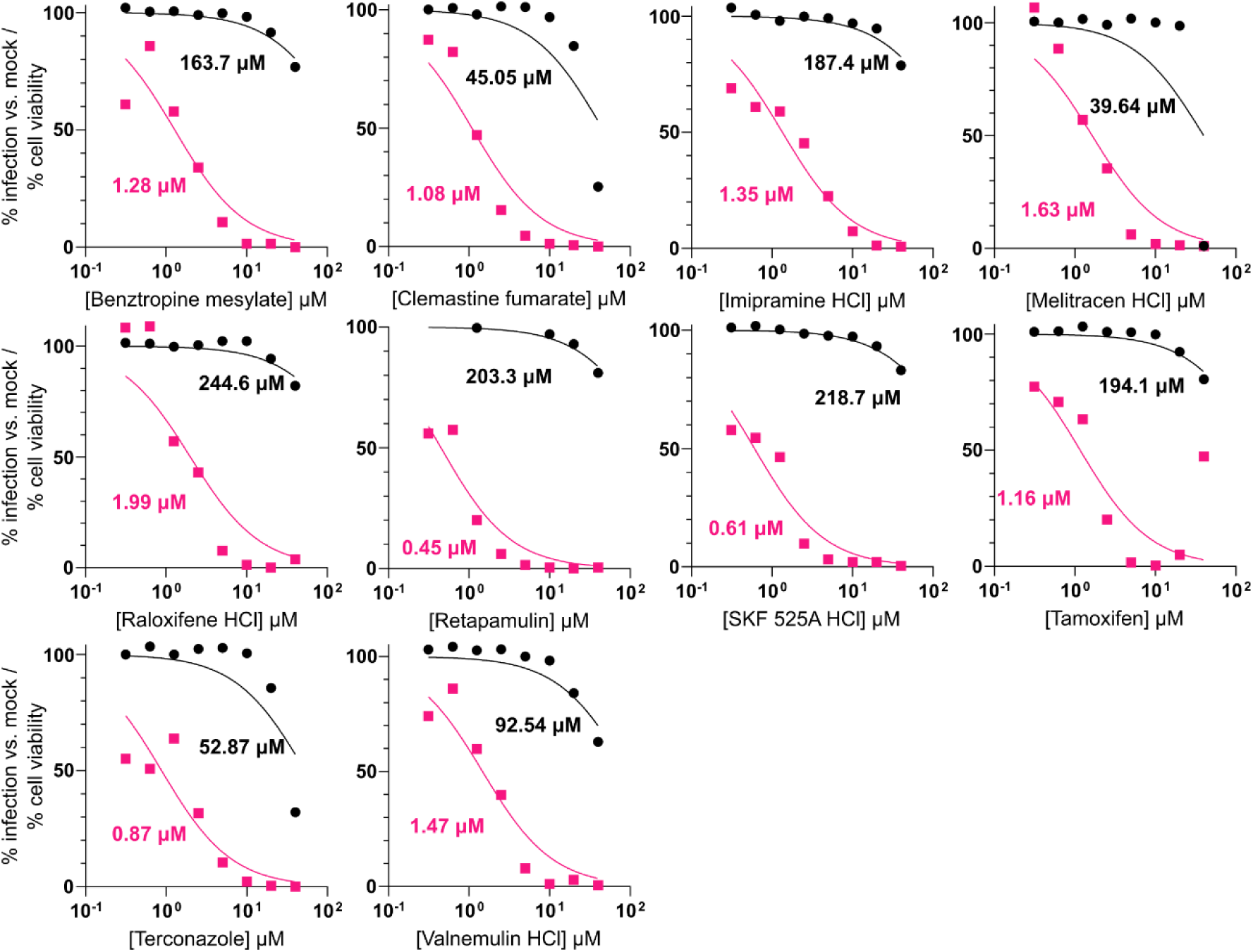
Dose-response curves of selected compounds (1). Infection levels are expressed as percentage relative to mock-treated controls. Pink curves represent EBOV pseudotype infection and black curves represent cell viability under the same treatment conditions. Half-maximal inhibition concentrations (IC_50_) or half-maximal cell cytotoxicity (CC_50_) are indicated for each virus-compound pair.

**Figure S3.**
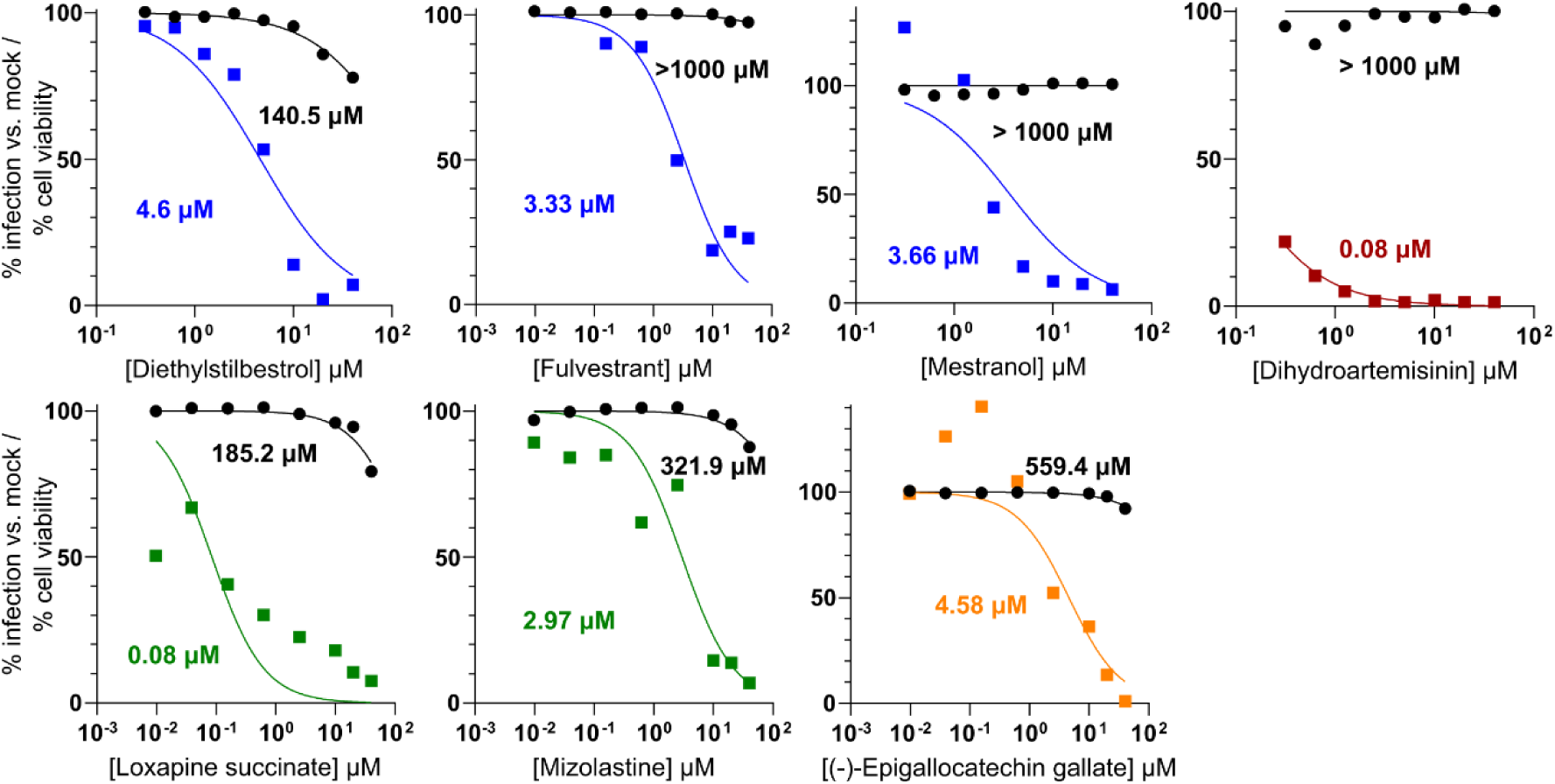
Dose-response curves selected compounds (bis). Infection levels are expressed as percentage relative to mock-treated controls. Curves correspond to the following pseudotypes: JUNV (blue), THOV (green), LPHV (orange), SINV (burgundy). Black curves represent cell viability under the same treatment conditions. Half-maximal inhibition concentrations (IC_50_) or half-maximal cell cytotoxicity (CC_50_) are indicated for each virus-compound pair.

**Figure S4.**
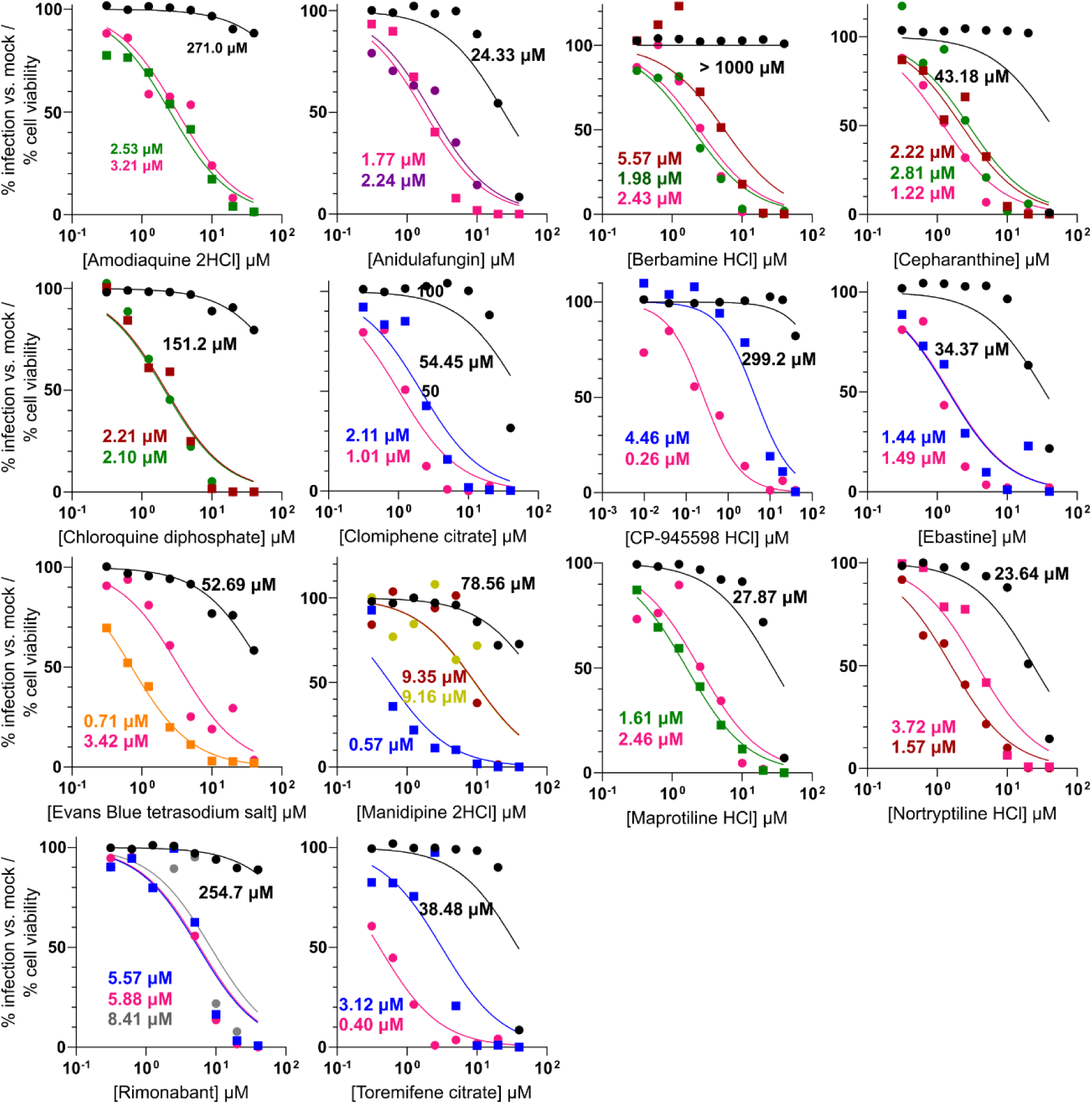
Dose-response curves of selected compounds (tris). Infection levels are expressed as percentage relative to mock-treated controls. Curves correspond to the following pseudotypes: EBOV (pink), JUNV (blue), THOV (green), LPHV (orange), SINV (burgundy), SNV (purple), RABV (yellow), and LACV (gray). Black curves represent cell viability under the same treatment conditions. Half-maximal inhibition concentrations (IC_50_) or half-maximal cell cytotoxicity (CC_50_) are indicated for each virus-compound pair.

**Figure S5.**
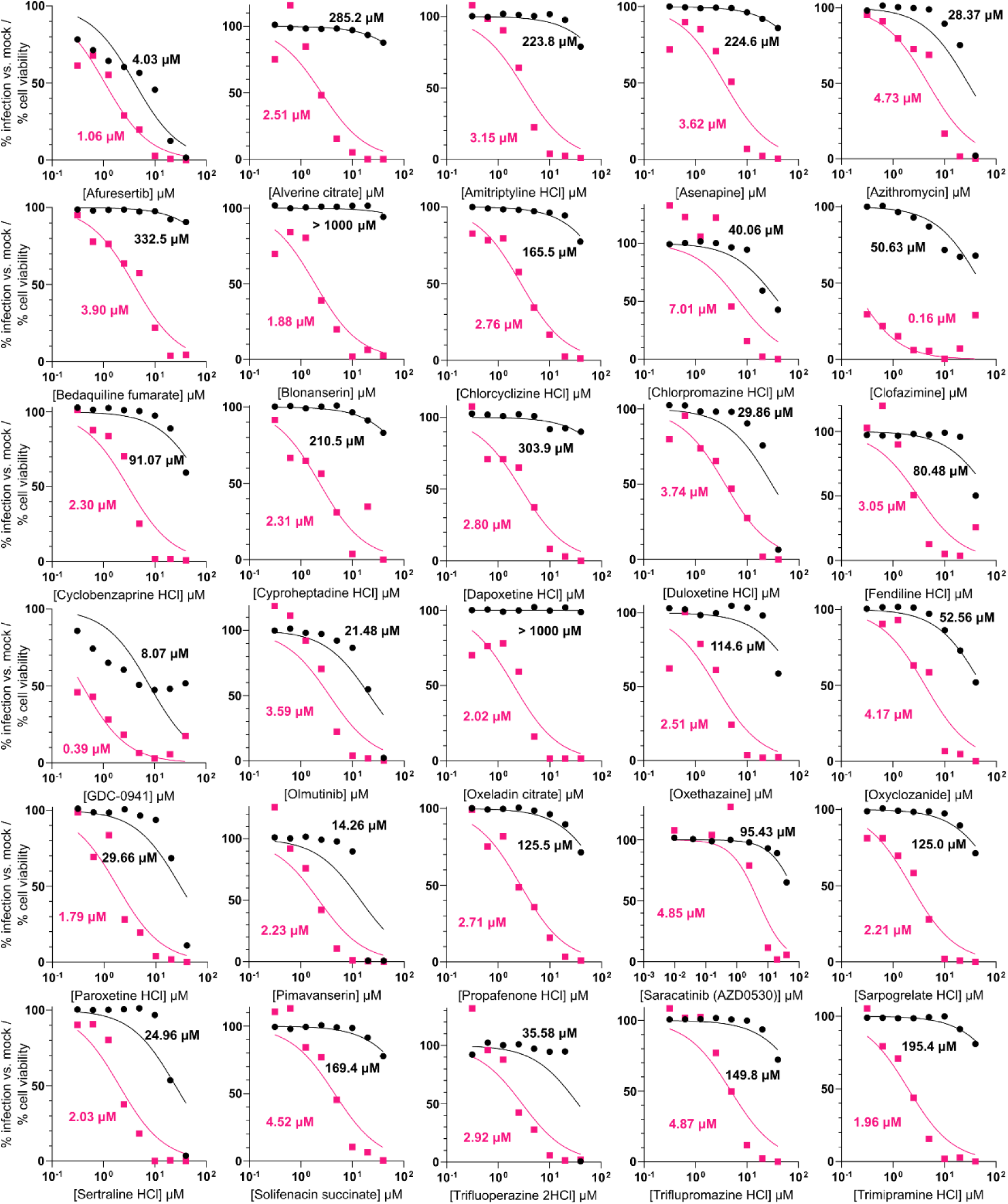
Dose-response curves of compounds against EBOV pseudotype not selected in Phase 2. Infection levels are expressed as percentage relative to mock-treated controls. Pink curves represent EBOV infection and black curves represent cell viability under the same treatment conditions. Half-maximal inhibition concentrations (IC_50_) or half-maximal cell cytotoxicity (CC_50_) are indicated for each virus-compound pair.

**Figure S6.**
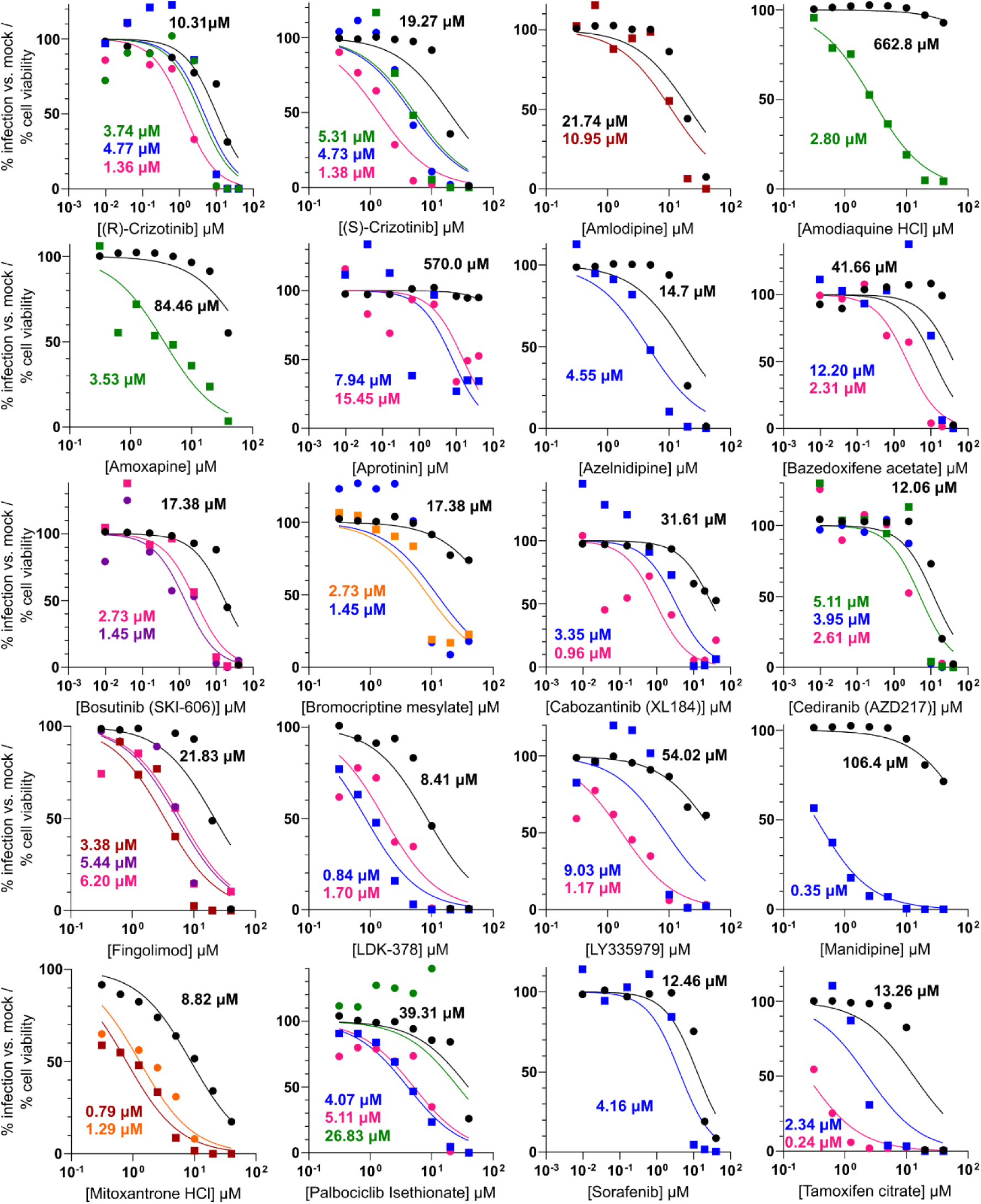
Dose-response curves of compounds not selected in Phase 2 (bis). Infection levels are expressed as percentage relative to mock-treated controls. Curves correspond to the following pseudotypes: EBOV (pink), JUNV (blue), THOV (green), LPHV (orange), SINV (burgundy), SNV (purple). Black curves represent cell viability under the same treatment conditions. Half-maximal inhibition concentrations (IC_50_) or half-maximal cell cytotoxicity (CC_50_) are indicated for each virus-compound pair.

**Figure S7.**
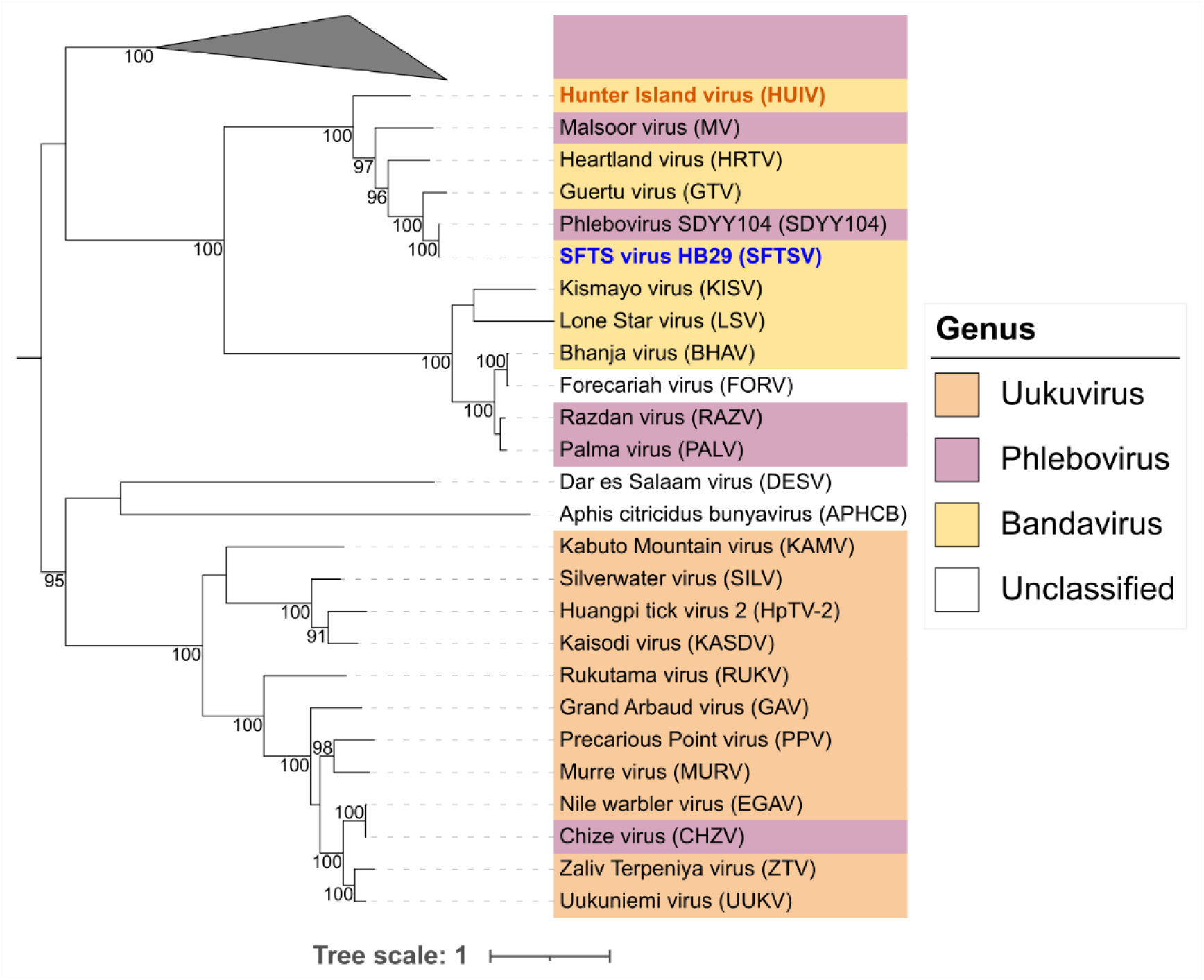
Phylogenetic tree of *Phenuiviridae* RBPs. Maximum-likelihood phylogenetic tree of RBPs from representative members of the *Phenuiviridae* family. Branch support values (bootstrap percentages) > 80 are indicated at key nodes. The blue name indicates the RBP initially included in the pseudotype panel, whereas the orange name denotes newly selected RBP. The scale bar represents amino acid substitutions per site.

**Figure S8.**
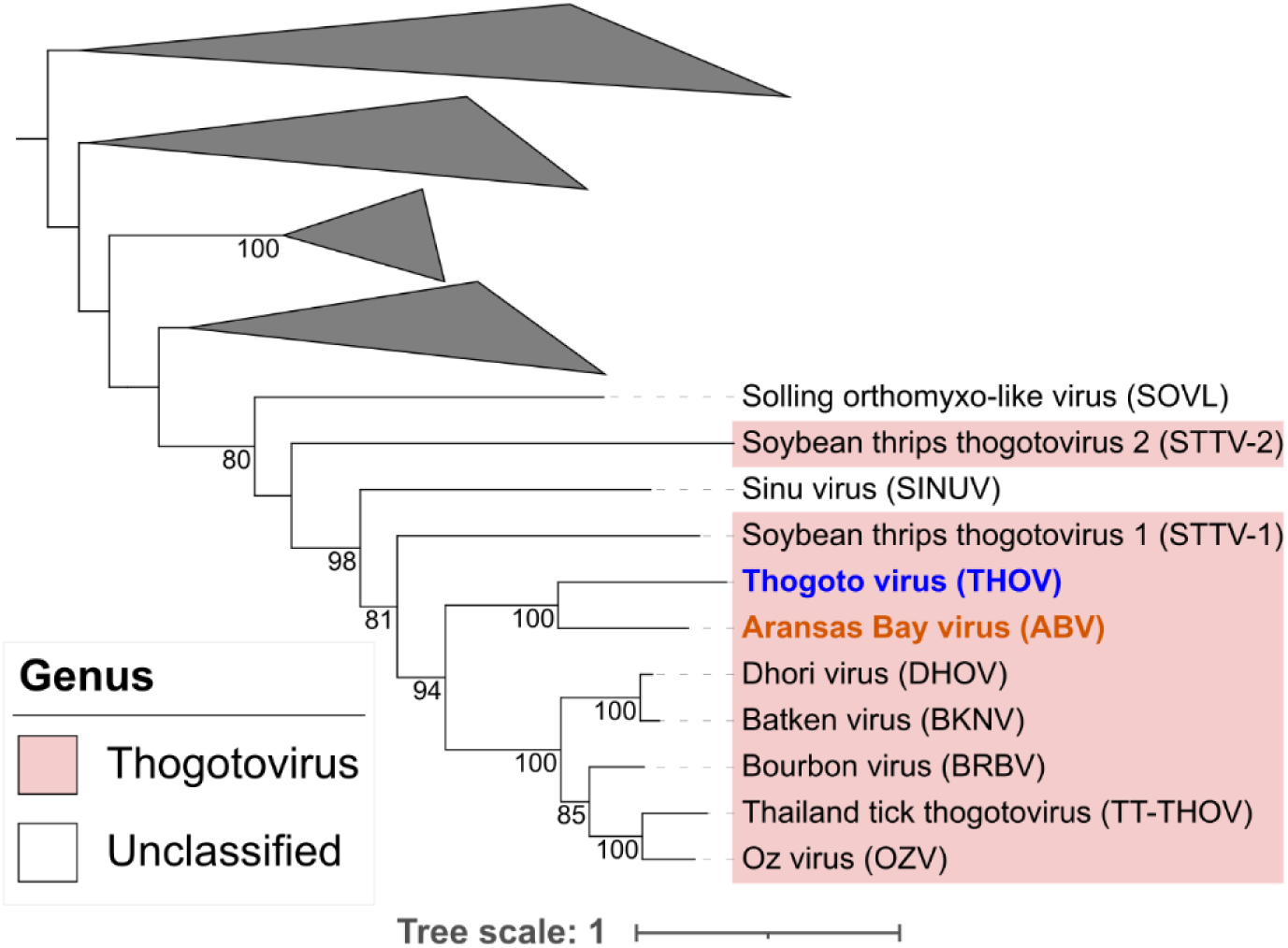
Phylogenetic tree of non-Influenza *Orthomyxoviridae* RBPs. Maximum-likelihood phylogenetic tree of RBPs from representative non-Influenza members of the *Orthomyxoviridae* family. Branch support values (bootstrap percentages) > 80 are indicated at key nodes. The blue name indicates the RBP initially included in the pseudotype panel, whereas the orange name denotes newly selected RBP. The scale bar represents amino acid substitutions per site.

**Figure S9.**
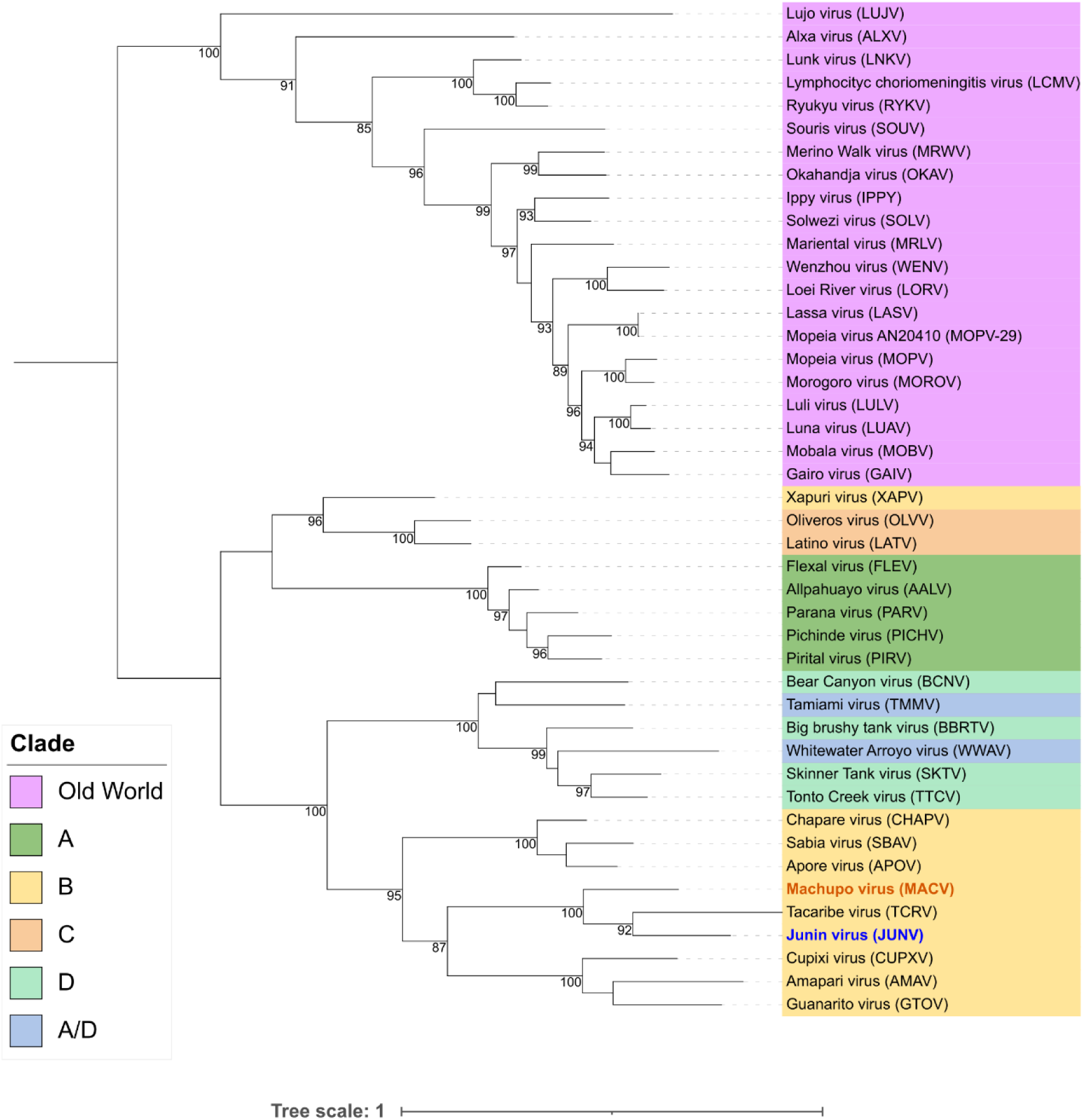
Phylogenetic tree of *Arenaviridae* RBPs. Maximum-likelihood phylogenetic tree of RBPs from representative members of the *Arenaviridae* family. Branch support values (bootstrap percentages) > 80 are indicated at key nodes. The blue name indicates the RBP initially included in the pseudotype panel, whereas the orange name denotes newly selected RBP.The scale bar represents amino acid substitutions per site.

**Figure S10.**
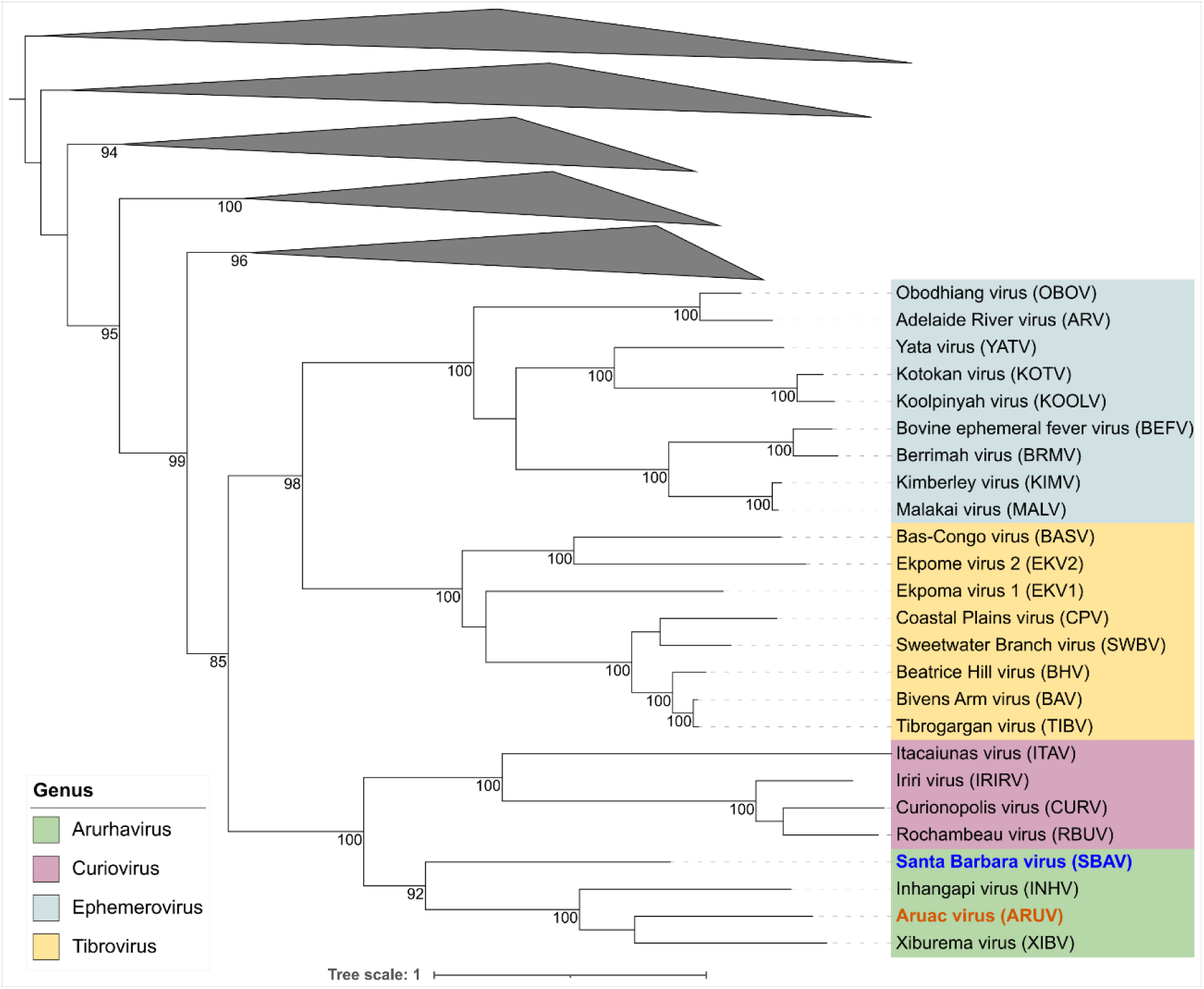
Phylogenetic tree of *Rhabdoviridae* RBPs. Maximum-likelihood phylogenetic tree of RBPs from representative members of the *Rhabdoviridae* family. Branch support values (bootstrap percentages) > 80 are indicated at key nodes. The blue name indicates the RBP initially included in the pseudotype panel, whereas the orange name denotes newly selected RBP.The scale bar represents amino acid substitutions per site.

**Figure S11.**
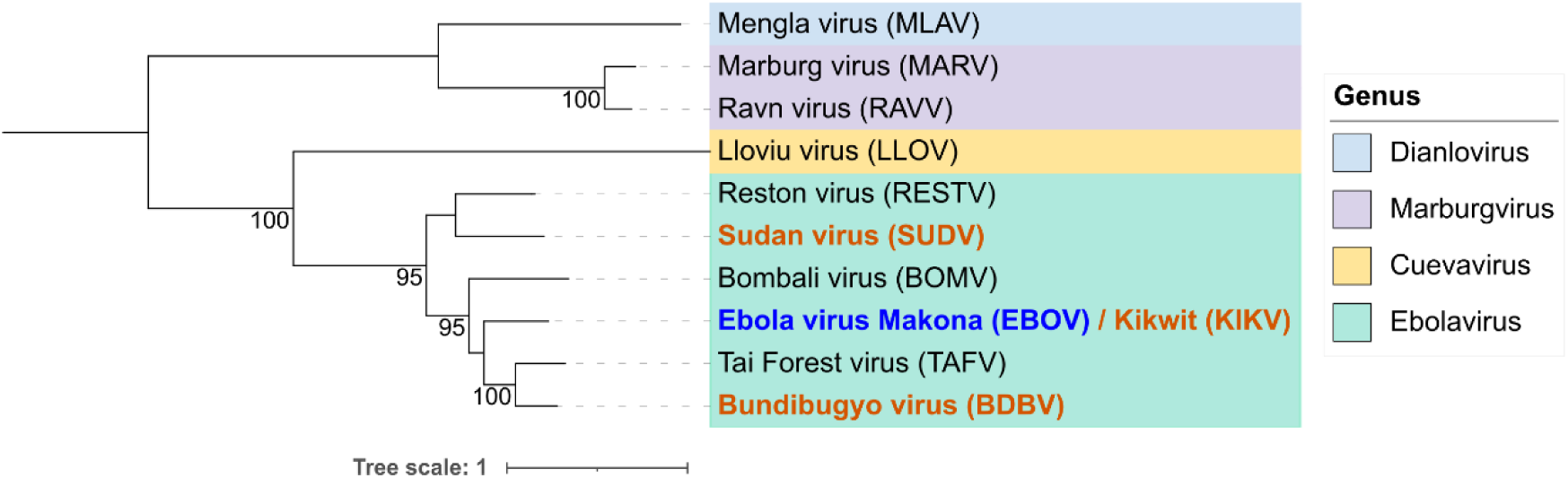
Phylogenetic tree of *Filoviridae* RBPs. Maximum-likelihood phylogenetic tree of RBPs from representative members of the *Filoviridae* family. Branch support values (bootstrap percentages) > 80 are indicated at key nodes. The blue name indicates the RBP initially included in the pseudotype panel, whereas the orange names denote newly selected RBP. The scale bar represents amino acid substitutions per site.

