## Supplementary Figures for "Anticipatory discovery of entry inhibitors against emerging viruses guided by viral phylogeny"

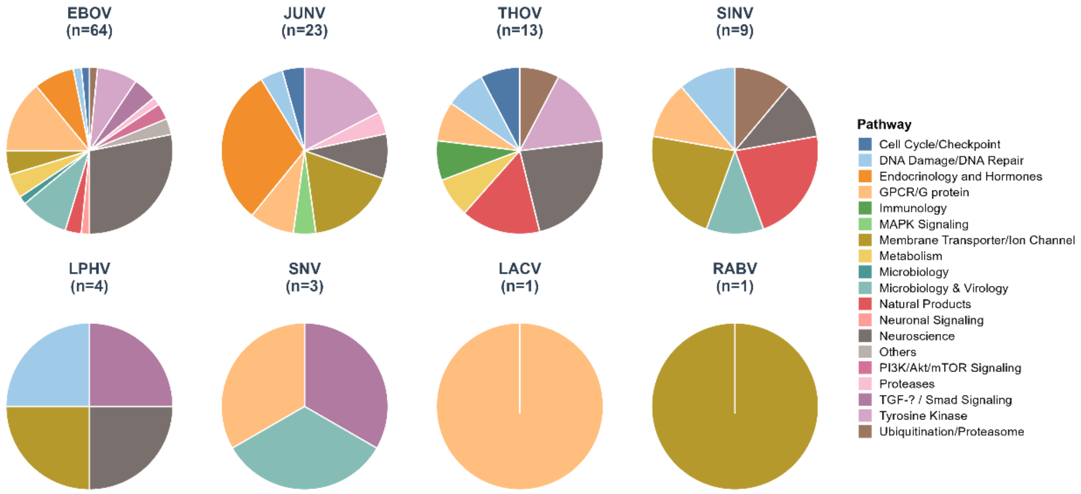


**Figure S1. Pathway distribution of hits identified for each RBP.** Pie charts represent the functional classification of compounds for each virus based on the annotation provided by APExBIO. For each viral pseudotype, the total number of selected hits is indicated in parentheses (n).

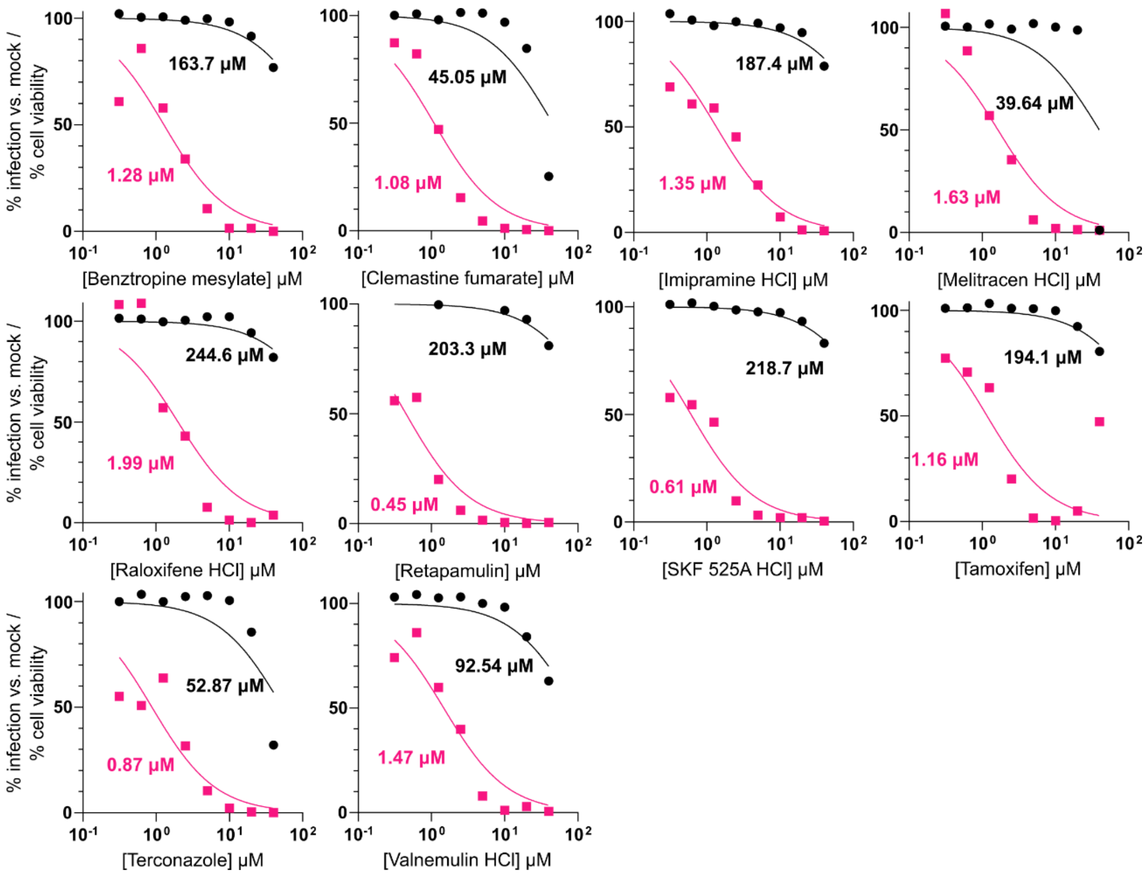


**Figure S2. Dose-response curves of selected compounds (1).** Infection levels are expressed as percentage relative to mock-treated controls. Pink curves represent EBOV pseudotype infection and black curves represent cell viability under the same treatment conditions. Half-maximal inhibition concentrations ($IC_{50}$) or half-maximal cell cytotoxicity ($CC_{50}$) are indicated for each virus-compound pair.

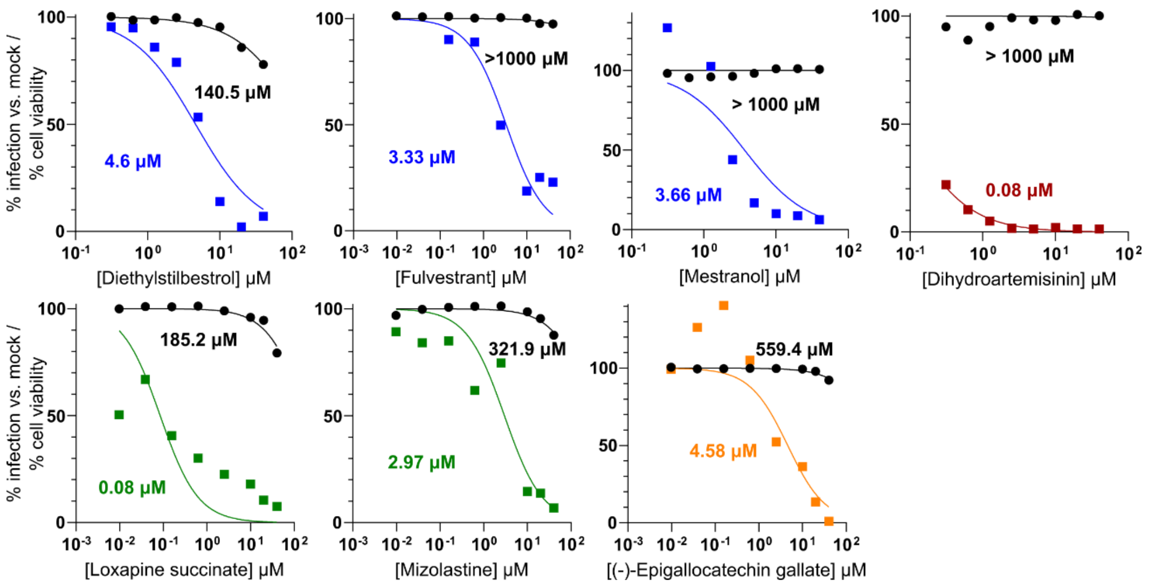


**Figure S3. Dose-response curves selected compounds (bis).** Infection levels are expressed as percentage relative to mock-treated controls. Curves correspond to the following pseudotypes: JUNV (blue), THOV (green), LPHV (orange), SINV (burgundy). Black curves represent cell viability under the same treatment conditions. Half-maximal inhibition concentrations ($IC_{50}$) or half-maximal cell cytotoxicity ($CC_{50}$) are indicated for each virus-compound pair.

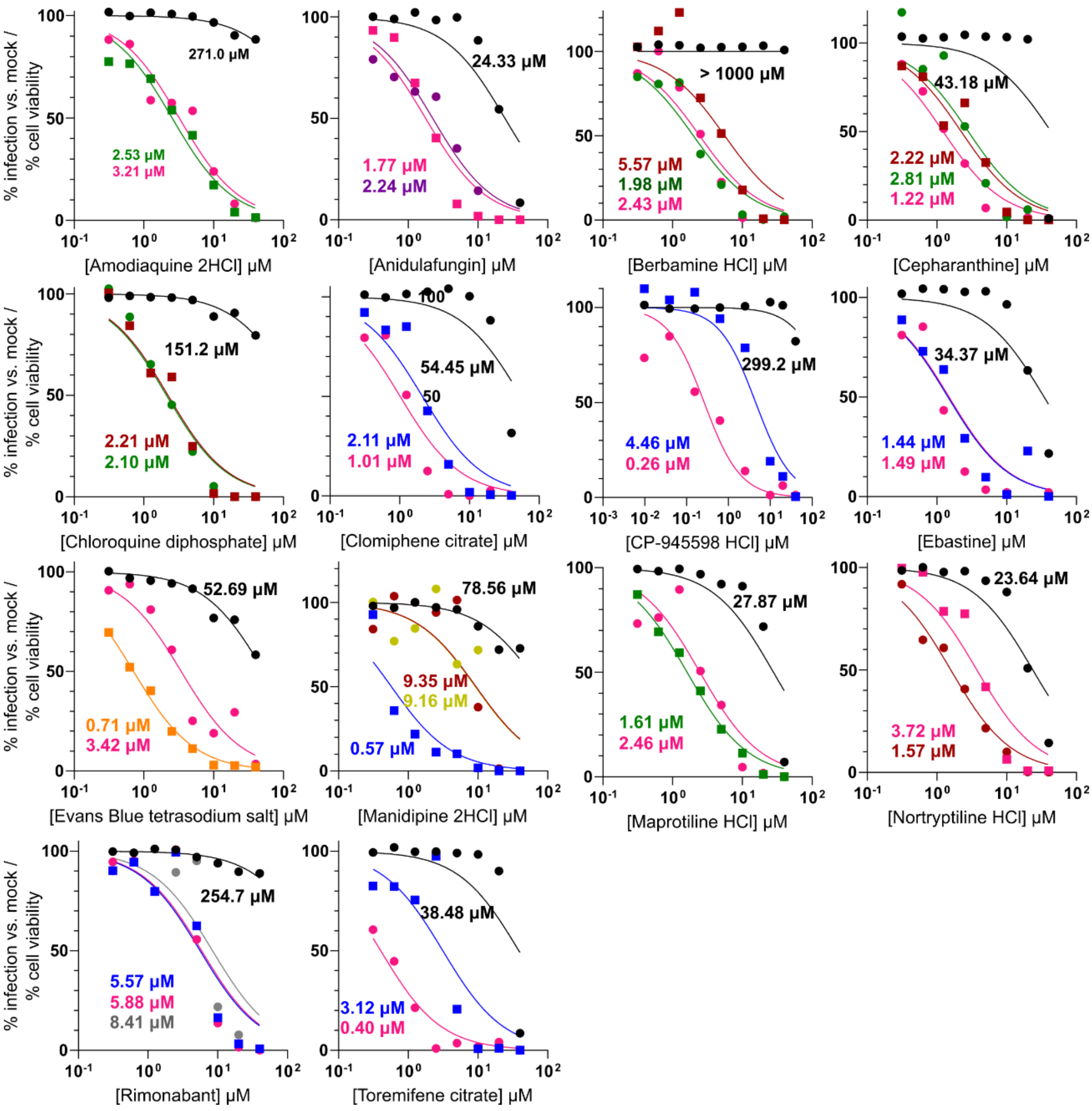


**Figure S4. Dose-response curves of selected compounds (tris).** Infection levels are expressed as percentage relative to mock-treated controls. Curves correspond to the following pseudotypes: EBOV (pink), JUNV (blue), THOV (green), LPHV (orange), SINV (burgundy), SNV (purple), RABV (yellow), and LACV (gray). Black curves represent cell viability under the same treatment conditions. Half-maximal inhibition concentrations ($IC_{50}$) or half-maximal cell cytotoxicity ($CC_{50}$) are indicated for each virus-compound pair.

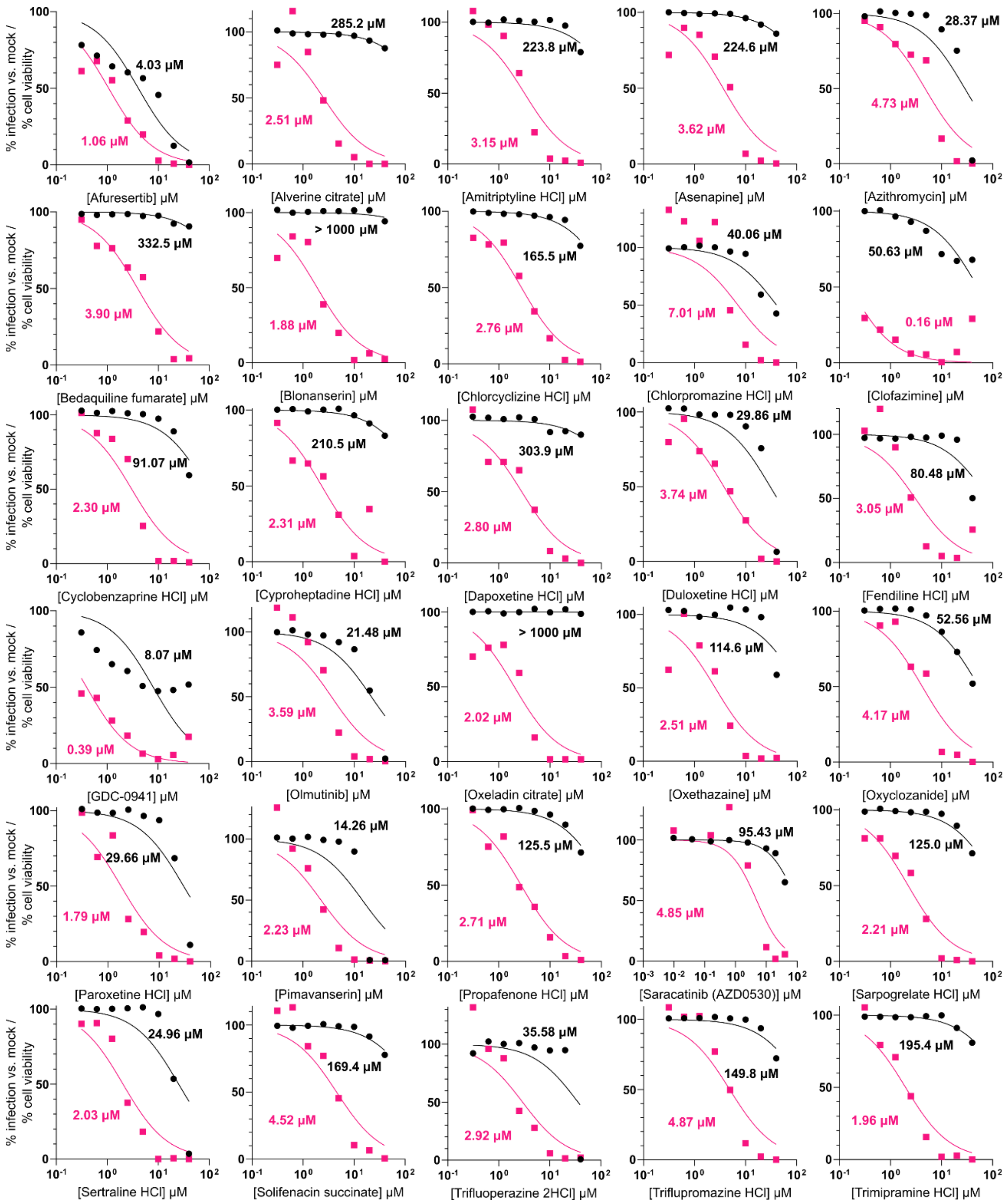


**Figure S5. Dose-response curves of compounds against EBOV pseudotype not selected in Phase 2.** Infection levels are expressed as percentage relative to mock-treated controls. Pink curves represent EBOV infection and black curves represent cell viability under the same treatment conditions. Half-maximal inhibition concentrations ($IC_{50}$) or half-maximal cell cytotoxicity ($CC_{50}$) are indicated for each virus-compound pair.

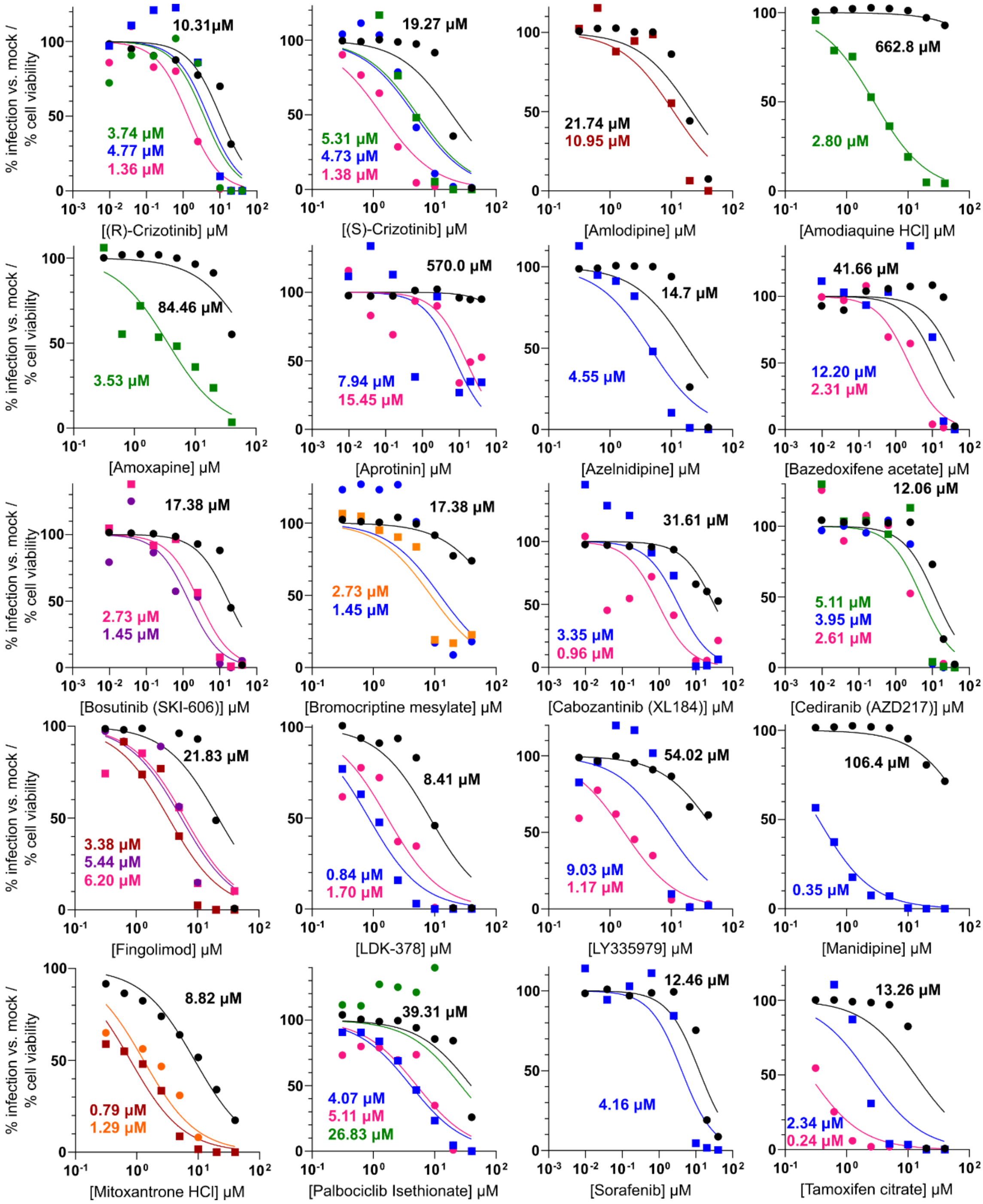


**Figure S6. Dose-response curves of compounds not selected in Phase 2 (bis).** Infection levels are expressed as percentage relative to mock-treated controls. Curves correspond to the following pseudotypes: EBOV (pink), JUNV (blue), THOV (green), LPHV (orange), SINV (burgundy), SNV (purple). Black curves represent cell viability under the same treatment conditions. Half-maximal inhibition concentrations ($IC_{50}$) or half-maximal cell cytotoxicity ($CC_{50}$) are indicated for each virus-compound pair.

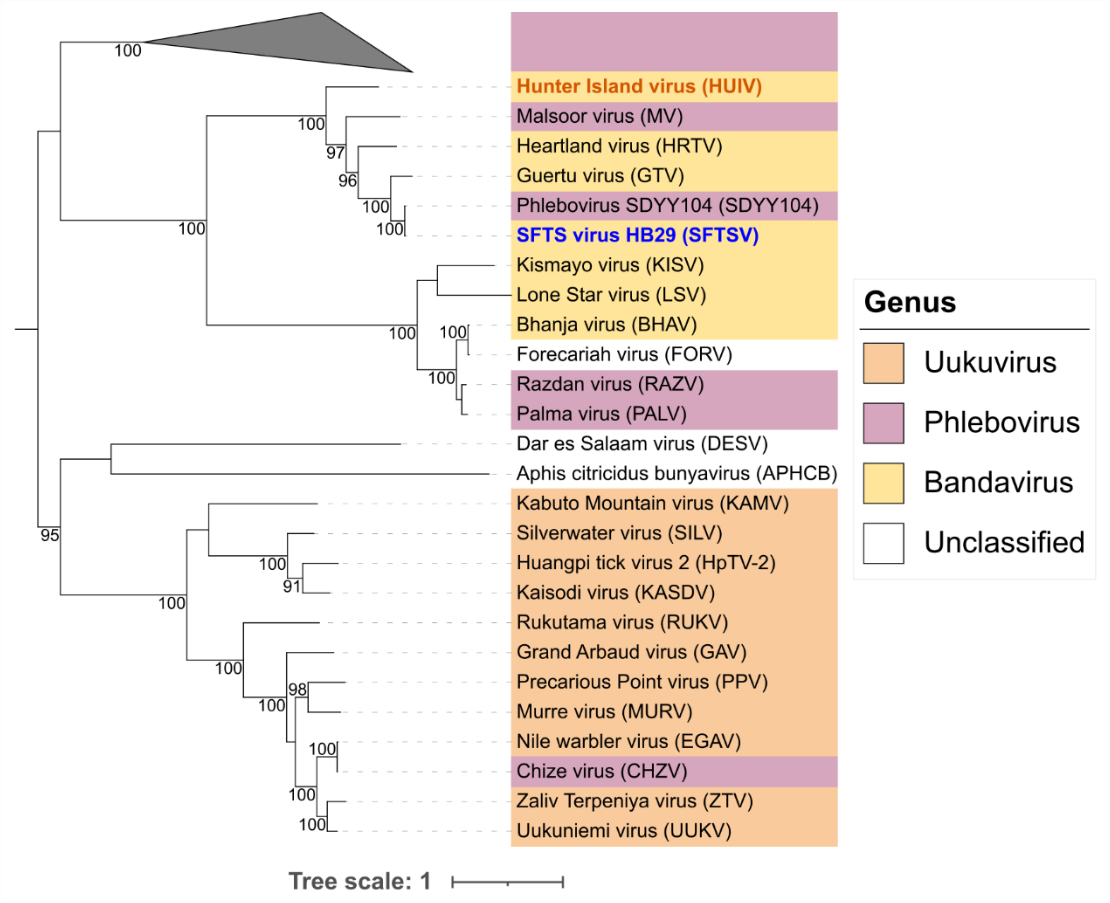


**Figure S7. Phylogenetic tree of *Phenuiviridae* RBPs.** Maximum-likelihood phylogenetic tree of RBPs from representative members of the *Phenuiviridae* family. Branch support values (bootstrap percentages) > 80 are indicated at key nodes. The blue name indicates the RBP initially included in the pseudotype panel, whereas the orange name denotes newly selected RBP. The scale bar represents amino acid substitutions per site.

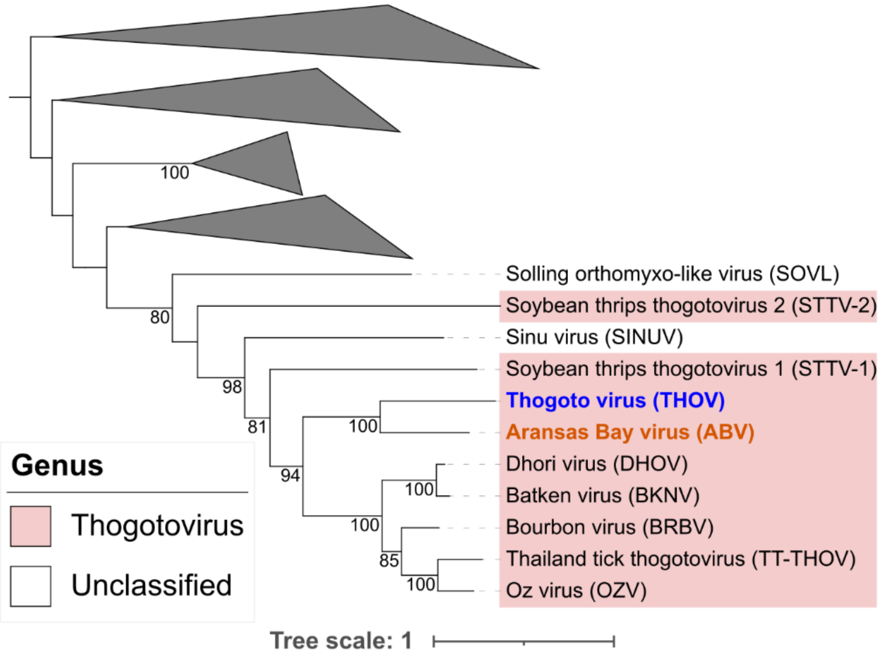


**Figure S8. Phylogenetic tree of non-Influenza *Orthomyxoviridae* RBPs.** Maximum-likelihood phylogenetic tree of RBPs from representative non-Influenza members of the *Orthomyxoviridae* family. Branch support values (bootstrap percentages) > 80 are indicated at key nodes. The blue name indicates the RBP initially included in the pseudotype panel, whereas the orange name denotes newly selected RBP. The scale bar represents amino acid substitutions per site.

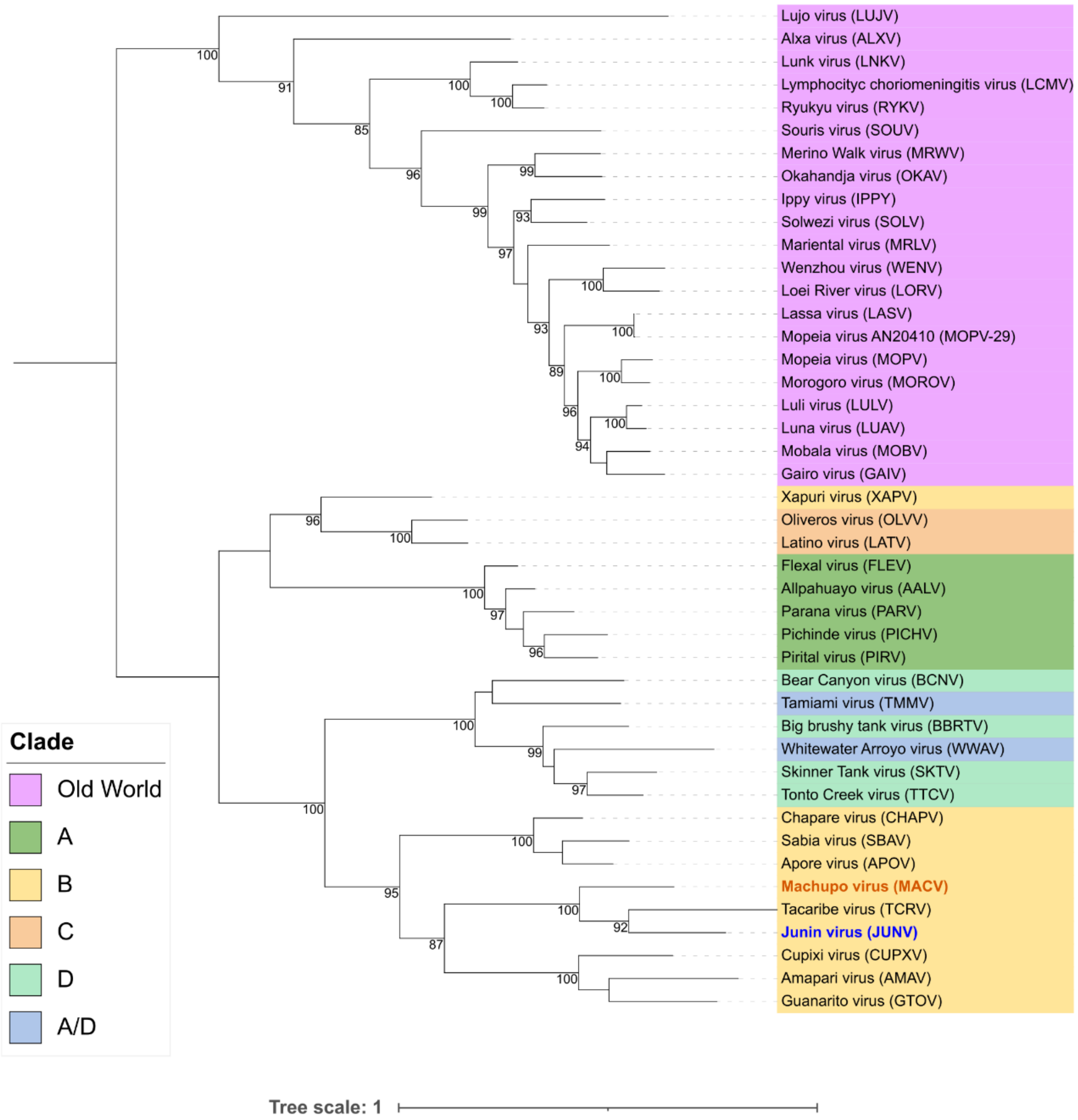


**Figure S9. Phylogenetic tree of *Arenaviridae* RBPs.** Maximum-likelihood phylogenetic tree of RBPs from representative members of the *Arenaviridae* family. Branch support values (bootstrap percentages) > 80 are indicated at key nodes. The blue name indicates the RBP initially included in the pseudotype panel, whereas the orange name denotes newly selected RBP.The scale bar represents amino acid substitutions per site.

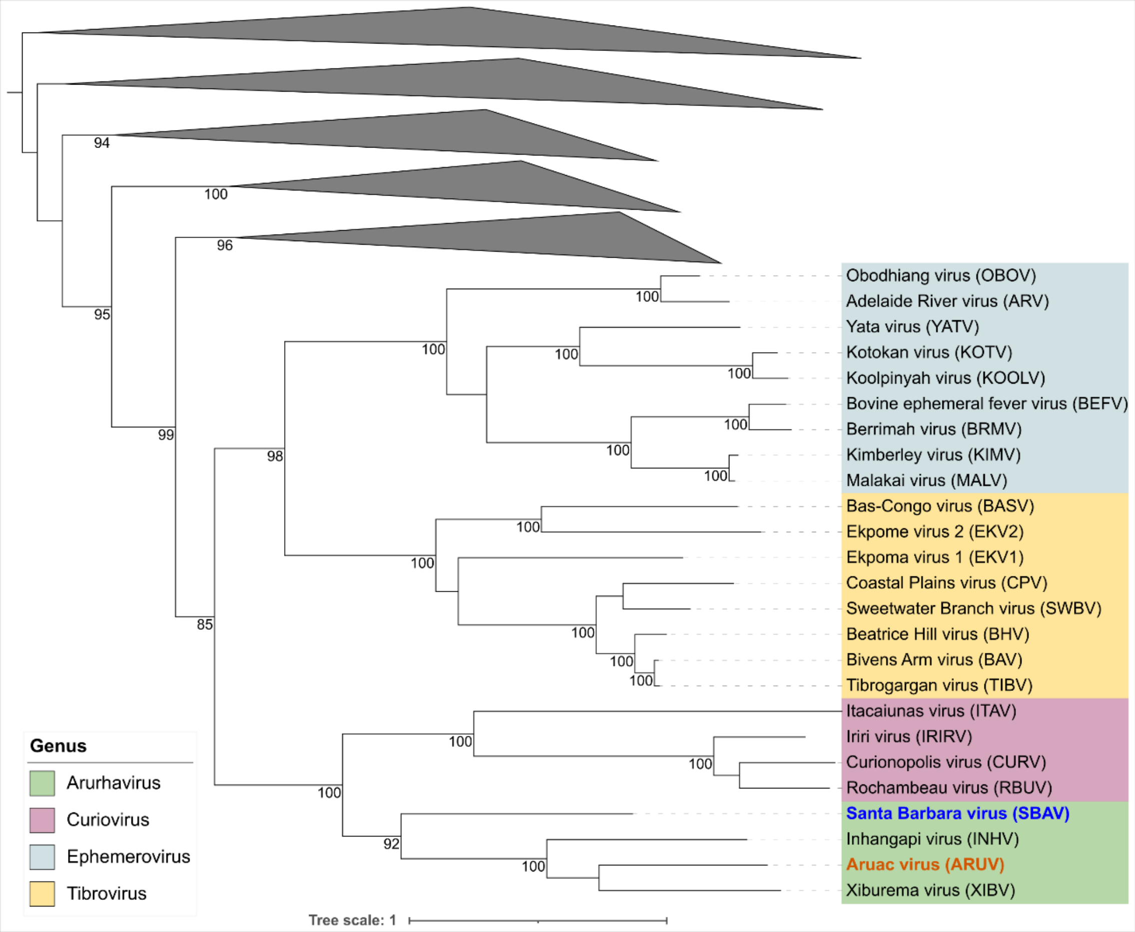


**Figure S10. Phylogenetic tree of *Rhabdoviridae* RBPs.** Maximum-likelihood phylogenetic tree of RBPs from representative members of the *Rhabdoviridae* family. Branch support values (bootstrap percentages) > 80 are indicated at key nodes. The blue name indicates the RBP initially included in the pseudotype panel, whereas the orange name denotes newly selected RBP.The scale bar represents amino acid substitutions per site.

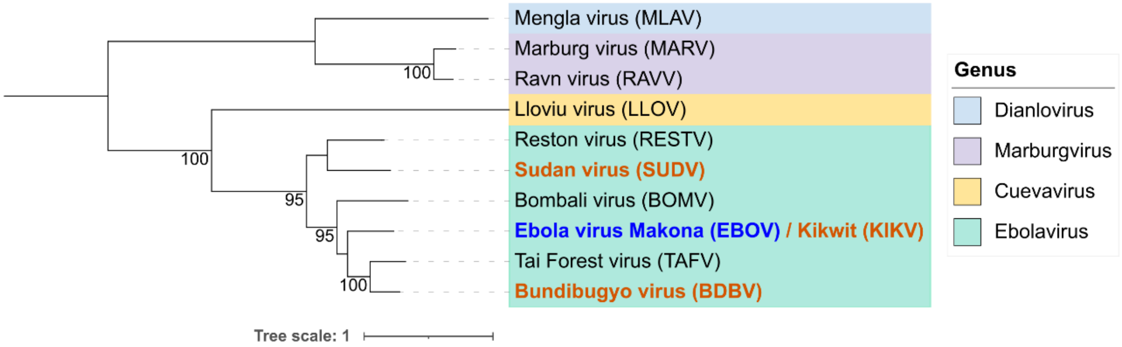


**Figure S11. Phylogenetic tree of *Filoviridae* RBPs.** Maximum-likelihood phylogenetic tree of RBPs from representative members of the *Filoviridae* family. Branch support values (bootstrap percentages) > 80 are indicated at key nodes. The blue name indicates the RBP initially included in the pseudotype panel, whereas the orange names denote newly selected RBP. The scale bar represents amino acid substitutions per site.
